# Dynamic Regulation of Vesicle Pools in a Detailed Spatial Model of the Complete Synaptic Vesicle Cycle

**DOI:** 10.1101/2023.08.03.551909

**Authors:** Andrew R. Gallimore, Iain Hepburn, Silvio Rizzoli, Erik De Schutter

## Abstract

Synaptic transmission is driven by a complex cycle of synaptic vesicle docking, release, and recycling, and maintained by distinct vesicle pools with differing propensities for docking and release. However, the partitioning of vesicles into distinct pools and the conditions under which they are recruited from the reserve to the recycling pool remain poorly understood. Whilst computational modeling at the molecular level has progressed massively in recent years, modeling voluminous, dynamic, and heterogenous structures such as synaptic vesicles remains an unsolved problem. Here, we use a novel vesicle modeling technology to model the complete synaptic vesicle cycle in unprecedented molecular and spatial detail at a hippocampal en passant synapse, incorporating vesicle diffusion, the accumulation and diffusion of proteins on the vesicle surface, vesicle clustering, tethering, and docking, and regulated vesicle fusion and recycling. Our model demonstrates highly dynamic and robust recycling of synaptic vesicles able to maintain stable and consistent synaptic release over time, even during high frequency and sustained firing. We also reveal how the cytosolic proteins synapsin-1 and tomosyn-1 can cooperate to regulate the recruitment of vesicles from the reserve pool during sustained periods of synaptic activity in order to maintain transmission, as well as the potential of selective vesicle tethering close to the active zone to ensure rapid vesicle replenishment and enhance the efficiency of the vesicle cycle by minimizing the recruitment of vesicles from the reserve pool.

## Introduction

Synaptic transmission is the fundamental mechanism of information transfer in the brain. Vesicles docked at the active zone rapidly fuse with the membrane in response to calcium influx following neural stimulation ^1–3^, driven by a complex signaling cascade involving membrane, vesicle, and cytosolic proteins ^4, 5^. These docked vesicles, primed for immediate release upon stimulation, form the readily-releasable pool (RRP) ^6, 7^. However, the fidelity of sustained transmission with repeated stimulation relies upon a reliable mechanism for the replenishment of the RRP as vesicles fuse with the membrane. The recycling pool – typically around 10-20% of the total vesicles – comprises freely-diffusing vesicles available to refill empty docking sites following vesicle fusion ^8–11^. A (generally much larger) pool of vesicles – accounting for 80-90% of the total vesicle population – known as the reserve pool, comprises vesicles resistant to release, intermixed with vesicles of the recycling pool ^11–14^. Pre-assembled vesicle proteins on the presynaptic membrane form an additional readily retrievable pool ^15, 16^. (Fig. 1a, b)

**1.**
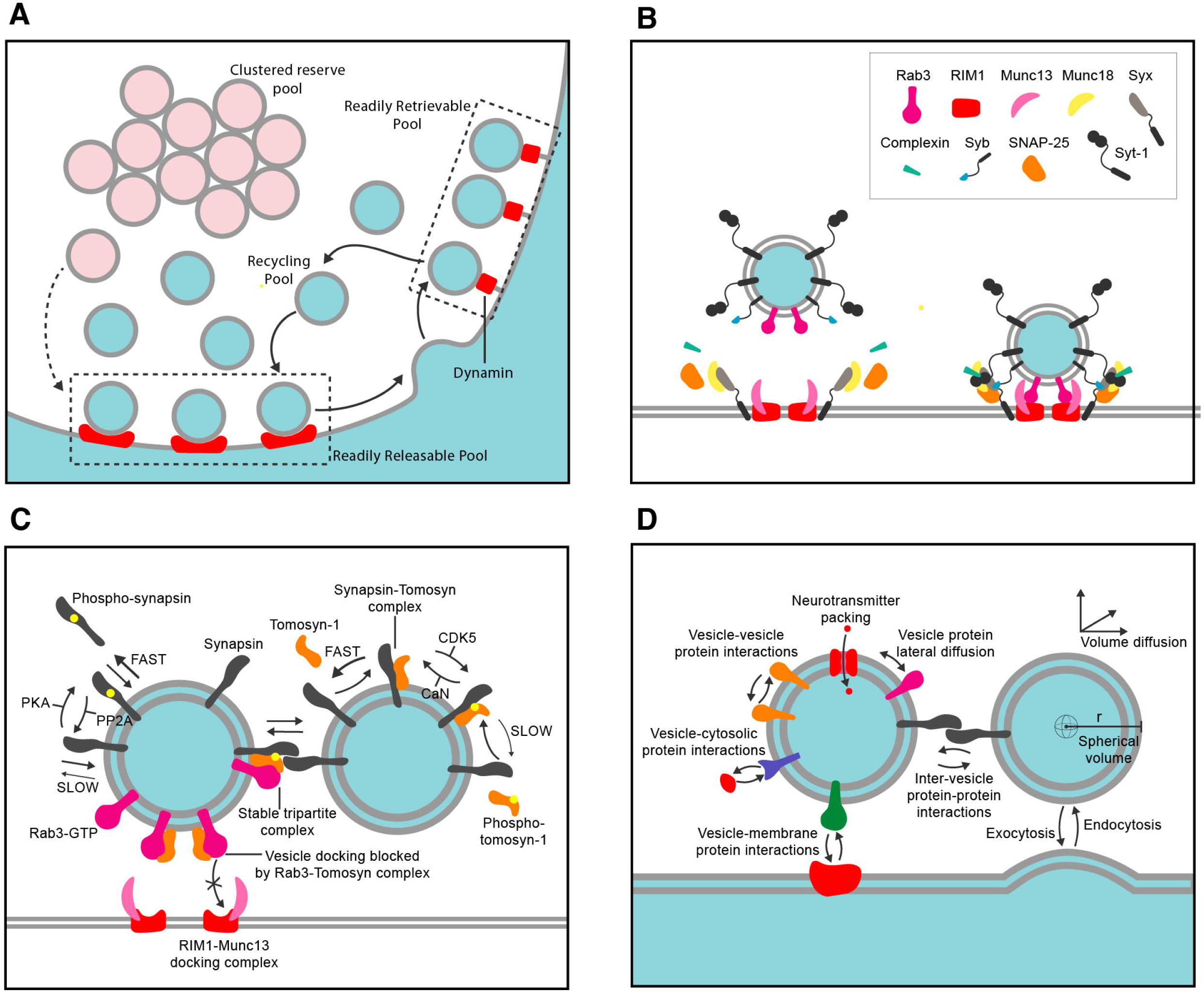
Structure of the vesicle and synaptic vesicle cycle model. A) Vesicle pools within the overall vesicle cycling model. B) Docking and priming interactions represented in the vesicle cycling model (see methods for details). C) Interactions between synapsin-1, tomosyn-1, and Rab3 that govern vesicle clustering and docking propensity. D) All behaviors and interactions of our model vesicles.

A typical hippocampal synaptic bouton contains 200-400 vesicles ^17–19^, which are maintained by inter-vesicle interactions between synapsin proteins on the vesicle surface, causing the reserve vesicles to form a cluster ^20–26^. Following endocytosis, a vesicle becomes part of the recycling pool, but slowly matures (over minutes to hours) and enters the reserve pool by becoming immobilized within the vesicle cluster by accumulation of synapsin ^9, 27, 28^. Synaptic activity causes synapsin dissociation from vesicles, controlled by PKA (and CaMKII) phosphorylation ^29^ dispersing the cluster and mobilizing vesicles for release ^30^.

Whilst the vesicle cluster plays an important role in maintaining vesicles within the bouton and restricting the availability of reserve vesicles for docking and release, the ability of vesicles to dock at the active zone is also modulated by the cytosolic protein tomosyn-1, which forms a complex with the small G-protein, Rab3-GTP, blocking its critical docking interaction with the active zone RIM1-Munc13 complex ^31, 32^. Notably, synapsin interacts in a phosphorylation state-dependent manner with the Rab3-GTP-tomosyn-1 complex to form a tripartite complex. CDK5 phosphorylates tomosyn-1 and enhances its interaction with synapsin-1 – this is counterbalanced by calcineurin (CaN)-dependent dephosphorylation ^33, 34^. This suggests a cooperative role of synapsin and tomosyn-1 in regulating the reserve pool using a combination of clustering and docking propensity to modulate the availability of vesicles for evoked release (Fig. 1c)

Although this regulated three-pool model is well accepted, many details are lacking. The characteristics of vesicles within the recycling and reserve pools, how they are partitioned, and the conditions under which vesicles are recruited from the reserve to the recycling pool remain poorly understood. Despite the increasing importance of computational modeling in understanding the behavior of complex emergent subcellular processes, such as long-term potentiation and depression ^35–42^, and synaptic vesicle release ^43, 44^, there are no published models of the synaptic vesicle cycle that incorporate the molecular mechanisms of vesicle transport, clustering, docking, priming, fusion, and recycling in a realistic spatial system. This is largely owing to a lack of technologies for modeling the complexities of vesicle structure and function. Whilst voxel– and particle-based reaction-diffusion systems ^45–48^ are appropriate for modeling networks of simple molecules with negligible volume, they are completely unsuitable for modeling large, mobile, and heterogenous structures such as synaptic vesicles, since their volume and molecular complexity have important roles in their function and influence their spatial dynamics and molecular interactions.

Our unique vesicle modeling technology extends the stochastic reaction-diffusion software, STEPS ^45, 46^, and allows us to model all key aspects of vesicle structure and function, including: vesicle diffusion; the accumulation and diffusion of proteins on the vesicle surface; inter-vesicle protein-protein interactions (controlling vesicle clustering); vesicle-cytosolic protein-protein (and small molecule) interactions; vesicle-surface protein-protein interactions (controlling vesicle tethering and docking); regulated endocytosis (vesicle recycling) and exocytosis (vesicle fusion and neurotransmitter release) ^49^. Each model vesicle occupies an excluded diffusible spherical volume and unique position within the model tetrahedral mesh and, in addition to the external surface exposed to the cytosol, possesses an internal compartment for the packing and release of neurotransmitters and other small molecules (Fig. 1d). Using this technology, we were able to model all major phases of the synaptic vesicle cycle at unprecedented levels of molecular and spatial detail, from docking and priming to fusion and recycling, in a realistic synaptic bouton morphology reconstructed from electron micrographs (serial-sectioning transmission electron microscopy) of a cultured hippocampal pyramidal cell ^50^ (Fig. 2a). Inter-vesicle protein-protein interactions also successfully replicated the dynamic liquid phase behavior of the vesicle cluster ^51, 52^ (Fig. 2b and supplementary video V1).

**2.**
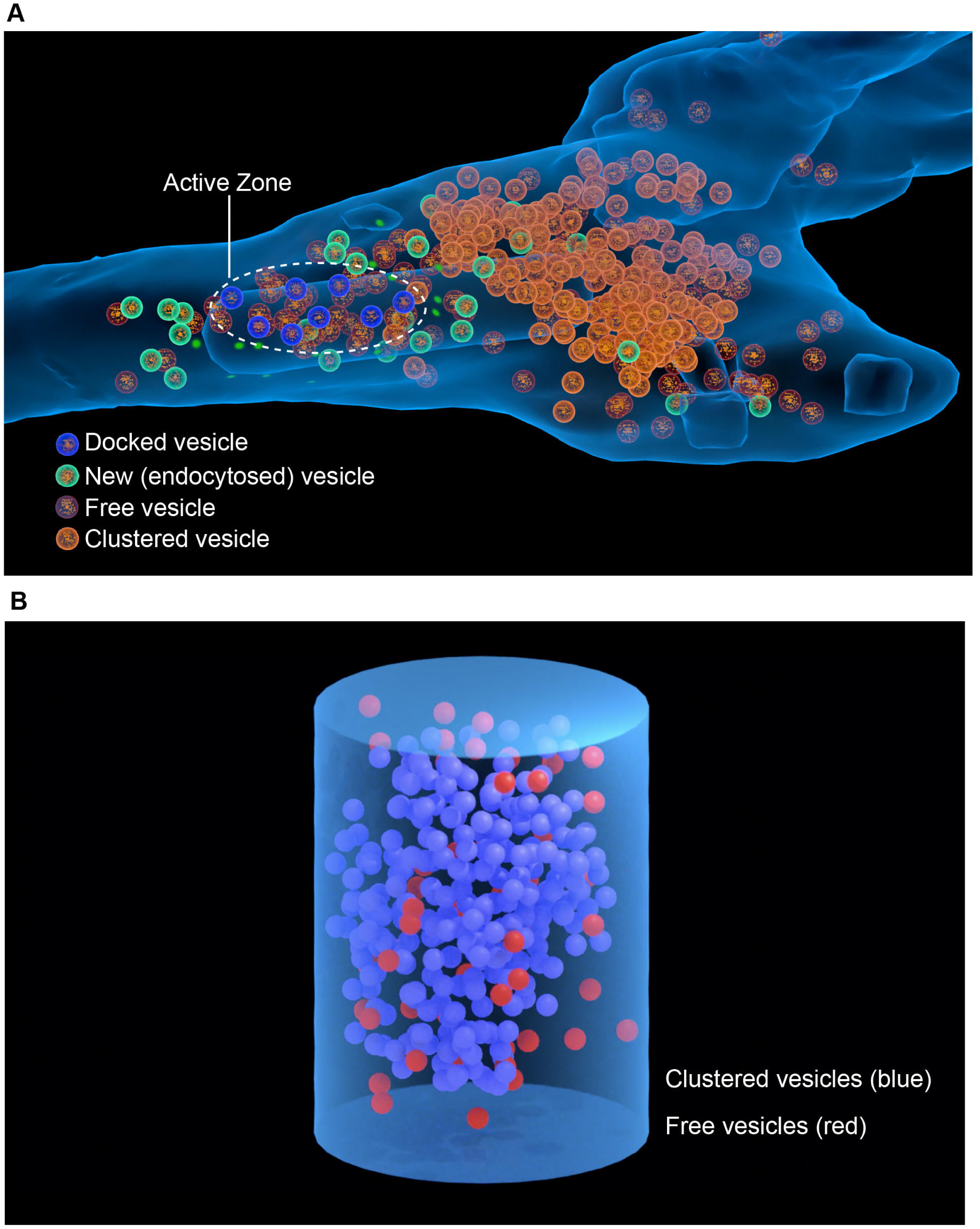
Blender visualization of the vesicle cycle model. A) Snapshot of complete vesicle cycling model showing clustered, free, newly endocytosed, and docked vesicles in the reconstructed synaptic bouton. B) Toy model in a cylindrical mesh showing vesicle clustering behavior (see supplementary video V1 for full animated visualization)

Our model reveals highly dynamic and robust recycling of synaptic vesicles able to maintain stable and consistent synaptic release over time, even during high frequency and sustained firing and assuming full vesicle collapse (as opposed to partial collapse proposed to occur during kiss and run recycling) followed by dispersal and retrieval of vesicle proteins. We also reveal how synapsin and tomosyn-1 can cooperate to regulate the recruitment of vesicles from the reserve pool during sustained periods of synaptic activity in order to maintain transmission, as well as the potential of selective vesicle tethering close to the active zone to ensure rapid vesicle replenishment and enhance the efficiency of the vesicle cycle by minimizing the recruitment of vesicles from the reserve pool.

## Results

### Release probability and recycling time

At hippocampal synapses, release probability (the probability of at least one vesicle fusion event following a single action potential) can vary greatly, with most synapses in the range of 0.1-0.6 ^53–55^ and largely dependent on the number of calcium channels coupled to each docking site ^55–57^. Our model, placing four calcium channels ^58^ in the vicinity of each docking site, displays a release probability of 0.5 and, as such, sits towards the higher end of this range. However, modeling a relatively high release probability synapse provides a better opportunity to model vesicle cycle dynamics within model simulation times that are feasible to compute.

The time taken for fused vesicles to be retrieved from the membrane (recycling time) varies depending on neuron type and is regulated at a cell-wide level ^59^, following an exponential distribution with a mean time, in hippocampal neurons, ranging from ∼5 to 40 seconds ^59, 60^. We measured the recycling time following 500 fusion events and observed recycling times between 2.5 and 18.6 seconds, with a mean time of 6.8 seconds and within the range observed experimentally (Supp. Fig. S2). Whilst our model doesn’t account for the time required for neurotransmitter refilling following vesicle endocytosis, proton gradient-driven neurotransmitter packing proceeds with similar kinetics to endocytosis (*τ≈*15s for glutamate via VGLUT) ^61, 62^ and, following vesicle retrieval, newly-retrieved vesicles have been shown to exhibit the same propensity for release as existing vesicles ^63^, suggesting that neurotransmitter packing is unlikely to be rate-limiting. Our model doesn’t exhibit very fast (<1 second) recycling events, which likely represent vesicle retrieval without the requirement for vesicle protein dispersal and re-accumulation or even without full vesicle collapse (“kiss and run”) ^64^ and, as such, is likely to be somewhat conservative compared to the real system.

### Endocytosis follows exocytosis closely, even at non-physiological firing rates

Following an initial 5 second equilibration time, we simulated neural firing for 45 seconds at frequencies ranging from 5 to 50 Hz (averaged over 20 simulations at each frequency) (Fig. 3). As expected, the rate of vesicle fusion increased with frequency: At 5, 10, and 20 Hz, an initial rapid fusion rate (corresponding to release of the RRP) was followed by a slower and almost linear rate of vesicle release. However, at 50 Hz, a further acceleration of release was observed after ∼15 seconds of stimulation (Fig. 3b). This effect could be blocked by selectively blocking tomosyn-1 dephosphorylation by CaN (Supp Fig. S3), demonstrating that this accelerated release is due to dephosphorylation of tomosyn-1, promoting dissociation of the synapsin-tomosyn-1 complex.

**3.**
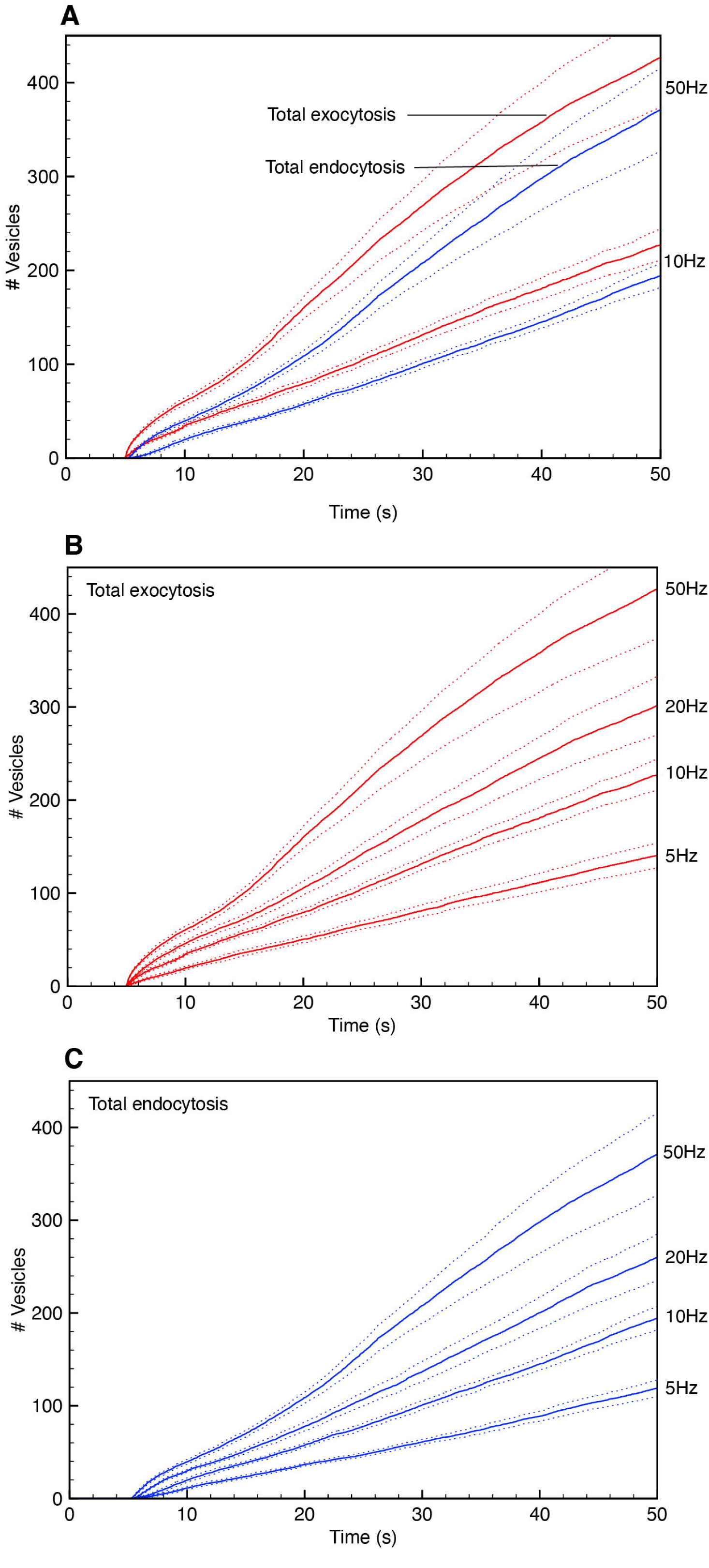
Synaptic vesicle release over time with frequency of stimulation. A) Total vesicle fusion and new vesicle endocytosis over time at 10Hz and 50Hz. (broken lines show 95% conf. interval) B) Total vesicle fusion over time at 5, 10, 20, and 50Hz. C) Total new vesicle endocytosis over time at 5, 10, 20, and 50Hz.

At all frequencies studied, endocytosis followed exocytosis closely with a delay of ∼2 seconds (Fig. 3a). The ratio of endocytosis:exocytosis remained between 0.84 and 0.87 at all frequencies from 5Hz to 50Hz ^65^. This suggests that, even with the assumption of dispersion and re-accumulation of vesicle proteins following exocytosis, the recycling system is capable of keeping pace with fusion at non-physiologically high firing rates.

### Vesicle material accumulates in the membrane with increasing frequency

The accumulation of vesicle material in the membrane was monitored by recording the number of collapsed vesicles (pits) over time. As the firing frequency increased, accumulation of vesicle material also increased, indicating that exocytosis was outpacing the rate at which vesicle material could be re-sequestered and vesicles recycled. This effect was particularly prominent at (non-physiological) 50Hz, with a 155% increase in vesicle material relative to baseline, compared to a 17% increase at 5Hz (Fig. 4a, c). This is comparable to experimental studies in hippocampal cells, in which accumulated membrane vesicle material was only observed to increase significantly with stimulation frequencies above 10Hz ^65^.

**4.**
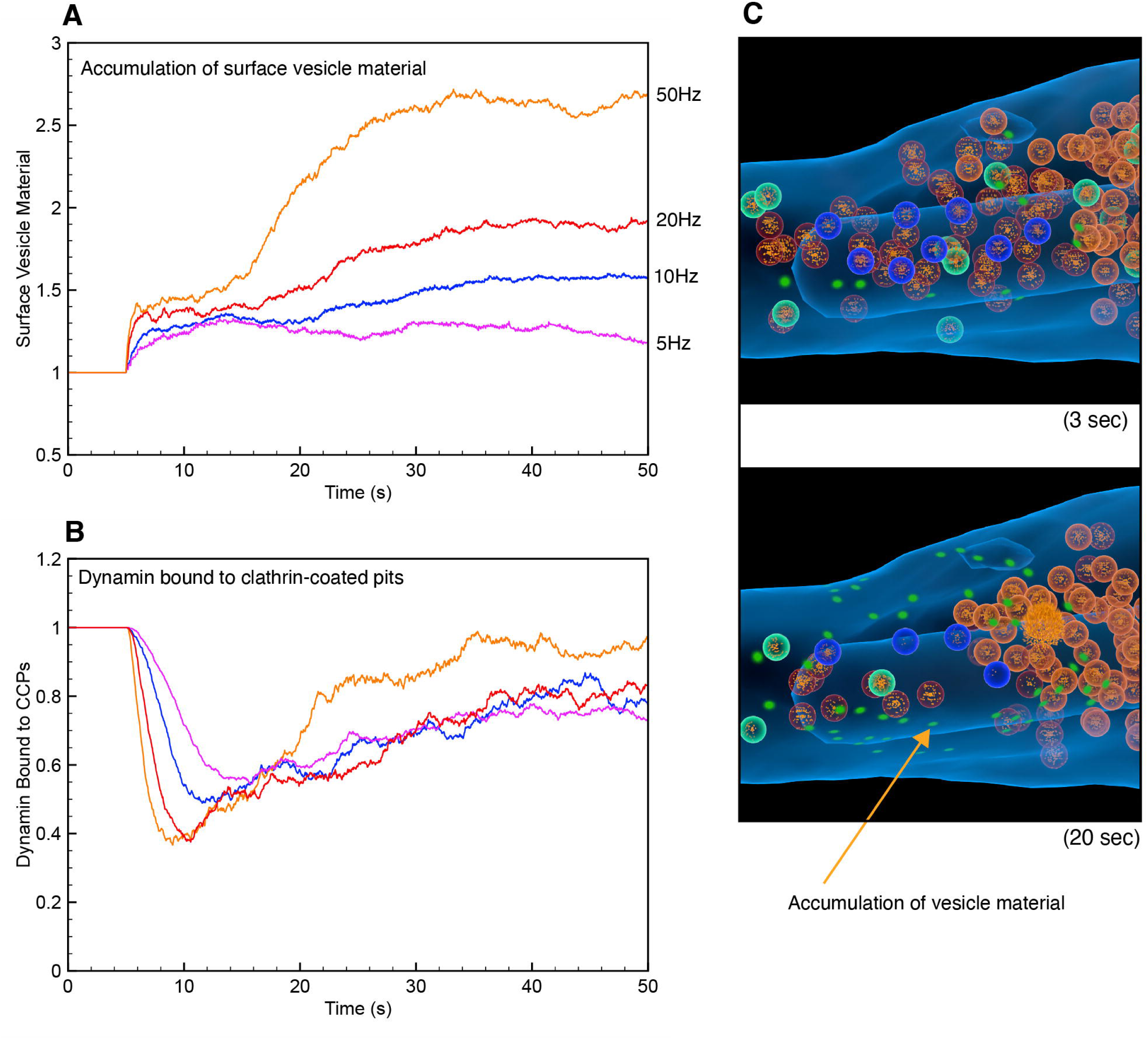
Accumulation of collapsed vesicle material in the plasma membrane. A) Accumulation of vesicle material in the membrane over time, at 5, 10, 20, and 50Hz stimulation. B) Dynamin bound to clathrin-coated pits (normalized to initial value) over time, at 5, 10, 20, and 50Hz. C) Accumulation of vesicle material is visible as green highlighted patches of membrane in the peri-active zone area.

### Dynamin Saturation with Firing Frequency

Dynamin-mediated scission is the final step in vesicle endocytosis ^60, 66, 67^ and, considering the dynamin copy number in a typical hippocampal bouton, a maximum of 50-60 fully-assembled dynamin complexes are available ^18^. As such, we monitored the total number of dynamin complexes in use (bound to a fully assembled clathrin-coated pit) over time. In the initial state, we assumed that each clathrin-coated pit was bound to a dynamin complex and thus primed for endocytosis. Immediately following stimulation, the number of bound dynamin complexes dropped to ∼40-60% of the initial number, indicating the endocytosis of vesicles from the readily-retrievable pool (Fig. 4b). This number slowly increased as fused vesicles accumulated in the membrane and entered the readily-retrievable pool. Whilst the rate of accumulation of bound dynamin increased with the frequency of stimulation, it never reached the saturation threshold, indicating that the number of dynamin complexes is not rate-limiting in maintaining endocytosis within the timeframe of the simulation.

### Reserve vesicle usage with frequency

In addition to monitoring the total number of vesicles exocytosed over time, we also separated fused vesicles into three types: vesicles from the initial recycling pool; vesicles from the initial reserve pool; and new vesicles formed by endocytosis. In experimental studies with cultured hippocampal neurons ^63^ ∼75% of the initial recycling pool was utilized following 300 action potentials, compared to only 230 action potentials on our model. However, this can be accounted for by the relatively high release probability of our model. At 10 Hz, reserve vesicles remained unused until ∼15s of stimulation and, at 45s, ∼20% of the reserve vesicles were utilized, compared to ∼98% of the initial recycling pool. The majority of vesicles used at 45s (∼65%) were newly formed (i.e. endocytosed), indicating that vesicle recycling is largely responsible for maintaining vesicle release over time (Fig. 5b). Reserve vesicle usage increased with frequency, with only 9% of reserve vesicles being utilized at 5Hz, increasing to 45% at 50Hz (Fig. 5b, c).

**5.**
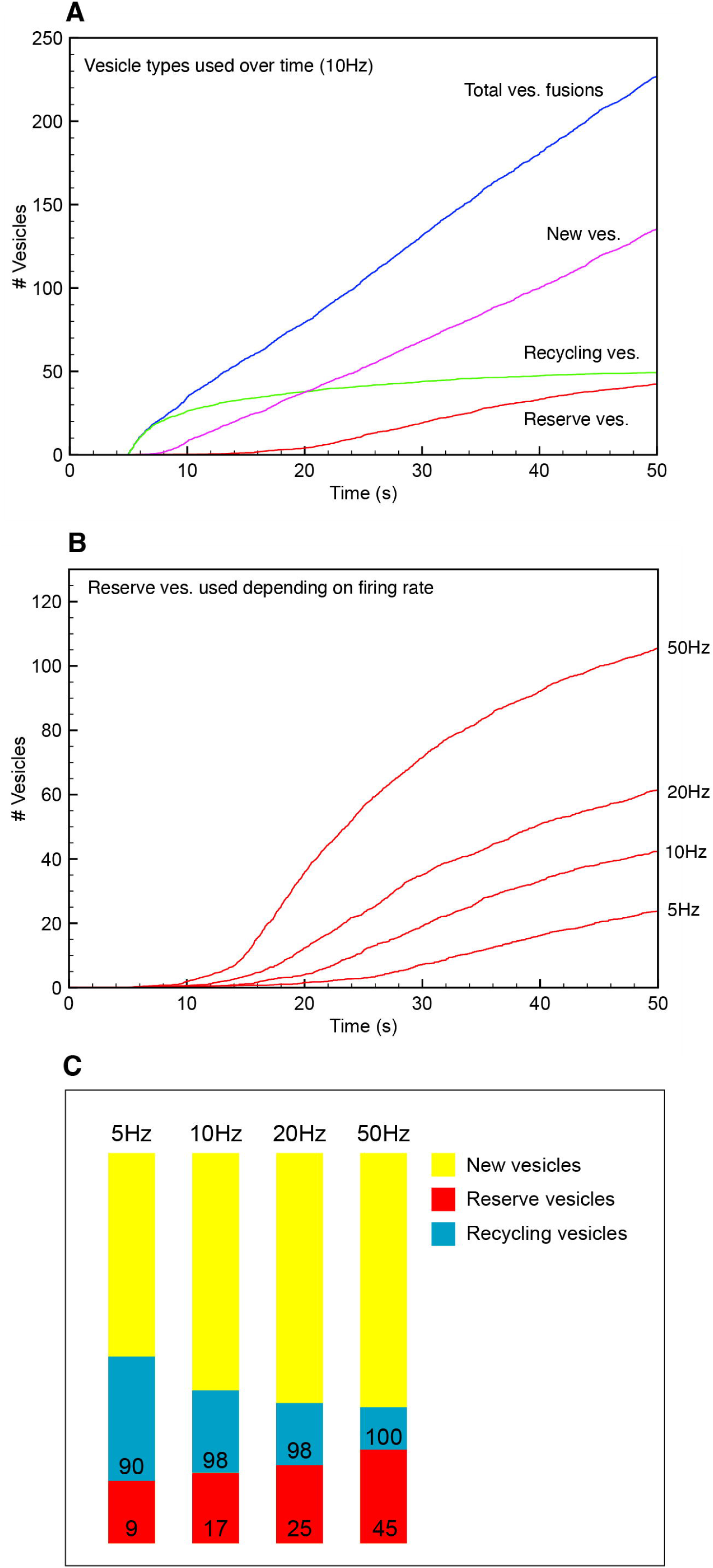
Usage of synaptic vesicles from different pools depending on frequency and PKA-mediated synapsin-1 phosphorylation. A) Fusions of different vesicle types during 10Hz stimulation. (reserve and recycling vesicles are from the initial population, new vesicles are all those formed by endocytosis). B) Proportion of different vesicle types used after 45 sec of stimulation at 5-50Hz. Numbers indicate percentage of initial population used after 45 sec. C) Usage of the reserve vesicle pool over time at 5, 10, 20, and 50Hz. D) The effect of blocking PKA-mediated phosphorylation of synapsin on total vesicle fusion events and reserve pool usage (10Hz).

### Dispersion of vesicle cluster is rapid and accompanied by a sharp increase in free tomosyn-1

The formation and dispersion of the vesicle cluster was monitored by counting the number of synapsin dimers between vesicles. Vesicles lacking any dimers, and thus not connected to any other vesicles, were counted as free, whereas those with dimer connections were considered part of the vesicle cluster. From the beginning of the simulation, the reserve vesicles became entirely clustered by ∼5s. The cluster remained completely intact with 7s of 10 Hz stimulation (Fig. 6b and supplementary video V2). The number of clustered reserve vesicles then declined to ∼60% after 24s of stimulation, before a rapid dissolution of the cluster over the following 3s. At 50Hz, cluster dispersion was complete by ∼17s of stimulation, driven by a faster rate of synapsin phosphorylation (Fig. 6a, d). Cluster dispersion was accompanied by the release of Tomosyn-1 from the vesicle surface, which depended on frequency, with up to ∼67% of tomosyn-1 being released from vesicles at 50Hz and ∼30% at 10Hz (Fig. 6b). This dispersion behavior is similar to that observed in frog motor terminals, in which vesicles remain immobile for ∼15s during 30Hz stimulation, before a sharp increase in mobility, indicating rapid dispersal of the vesicle cluster ^68^. Blocking PKA-dependent phosphorylation of synapsin, and thus dispersal of the vesicle cluster, blocked reserve vesicle usage completely and reduced overall vesicle usage by 17% (at 10Hz). (Fig 6c)

**6.**
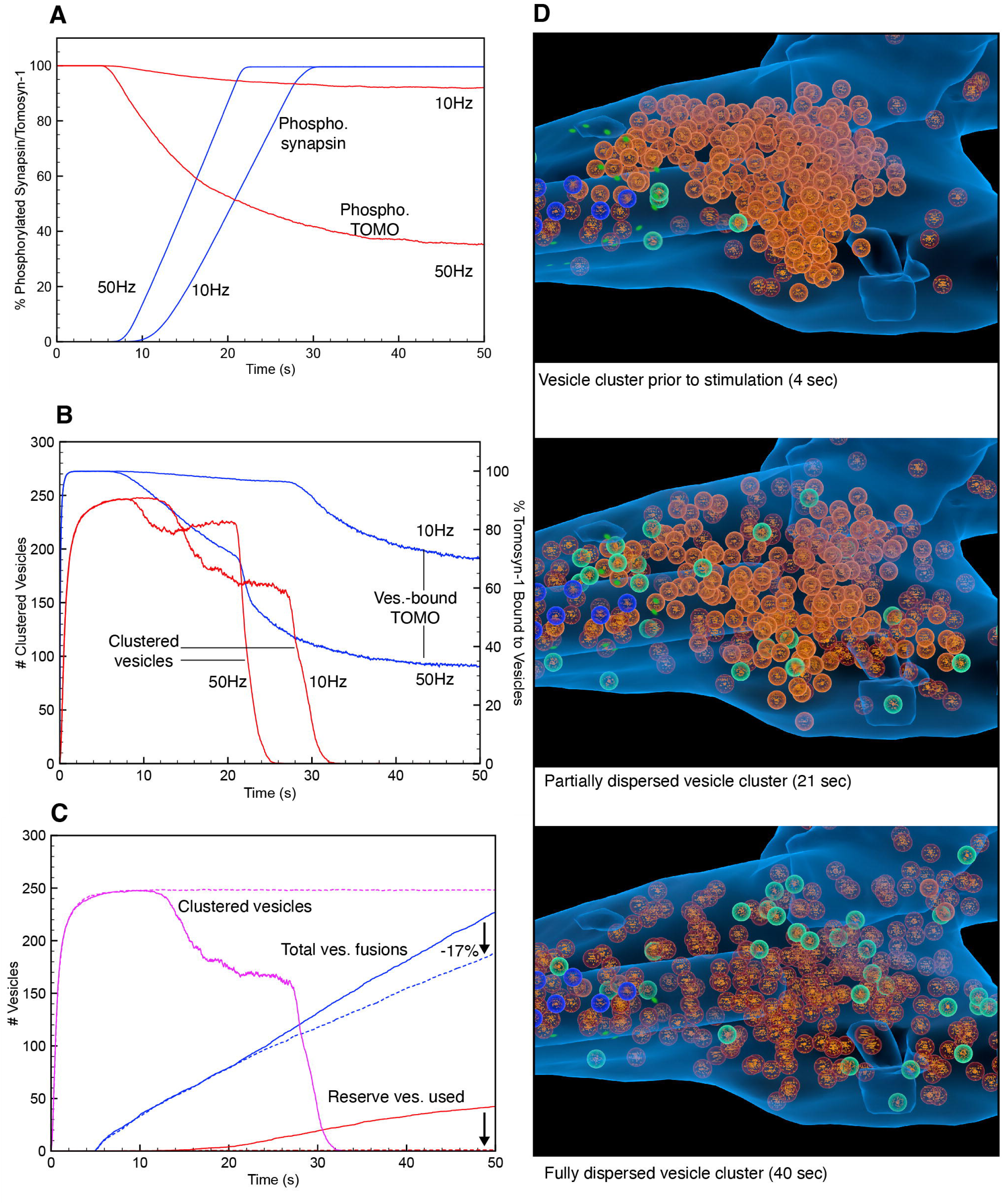
Regulation of vesicle clustering and dispersion by synapsin-1 and tomosyn-1. A) Phosphorylation of synapsin1 and tomosyn-1 over time at 10 and 50Hz stimulation over 45 sec. B) Vesicle cluster formation and dispersal and tomosyn-1 bound to vesicles over time at 10 and 50Hz stimulation beginning at 5 sec. C) Effect of blocking cluster dispersion (by blocking PKA-mediated synapsin phosphorylation) on vesicle fusion and reserve vesicle usage with 10Hz stimulation. D) Progression of vesicle cluster dispersal over time with 50Hz stimulation (see supp. video V2).

### Tomosyn-1 expression and vesicle clustering regulate the release of vesicles from the reserve pool

In hippocampal neuronal cultures, knockdown of tomosyn-1 causes a 30-50% increase in the apparent size of the total releasable pool of vesicles, whereas tomosyn-1 overexpression reduces this pool by ∼25% ^31^. We observed similar responses to tomosyn-1 knockout and doubling the tomosyn-1 copy number in our model. Relative to the control, the tomosyn-1-null model increased the total number of vesicles released from the initial recycling and reserve pools by 70% (Fig. 7a). However, this effect was entirely caused by an increase in reserve pool vesicle usage: The number of initial recycling pool vesicles used actually decreased by 18%. Interestingly, usage of newly endocytosed (recycled) vesicles decreased by 33% in the tomosyn-1-null condition, indicating that the vesicle cycle was relying much more heavily on reserve pool vesicles to maintain synaptic release. However, the total number of vesicles used overall increased by 8%, owing to the absence of tomosyn-1’s inhibitory effect on docking (Fig. 7b). Doubling the tomosyn-1 copy number had no significant effect on the rate of vesicle release up to ∼15s of stimulation at 10Hz. However, by strongly suppressing reserve pool vesicle usage and vesicle docking, fusion rate then slowed and the total vesicles used at 45 seconds was reduced by 29%. (Fig. 7c,d)

**7.**
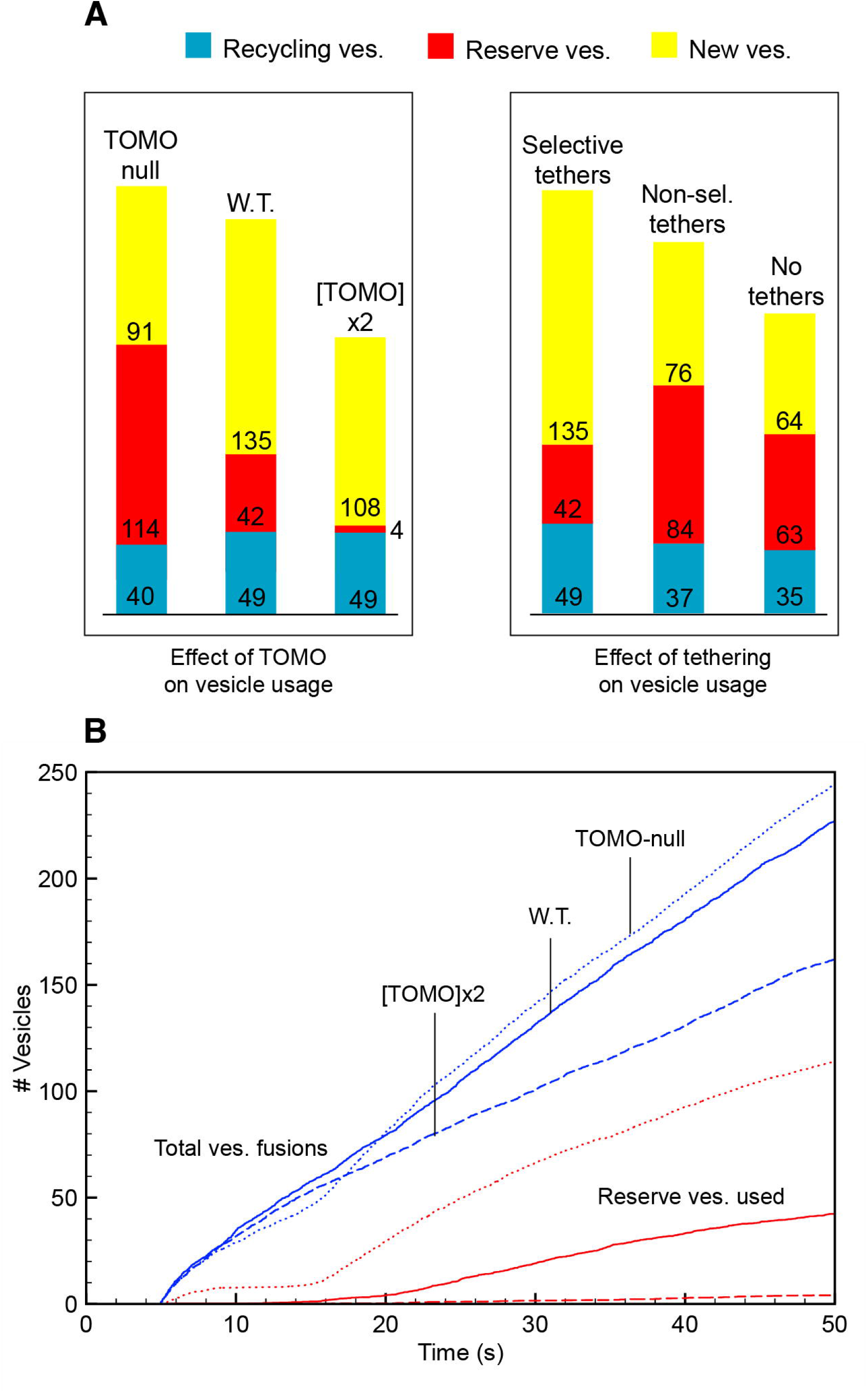
Effect of tomosyn-1 and tethering on synaptic vesicle usage. A) Effect of tomosyn-1 copy number, selective and non-selective tethering on vesicle usage after 45 sec of stimulation at 10hz. Left: Effect of tomosyn-1 copy number of absolute number of vesicles used from each pool. Numbers represent number of vesicles used after 45 sec of stimulation); Right: Effect of tethering (selective and non-selective) on absolute number of vesicle types used from each pool. B) Effect of tomosyn-1 (TOMO) copy number on total vesicle fusion and reserve vesicle usage over time at 10Hz.

Together, these results indicate that tomosyn-1 not only inhibits docking but, by forming a stable tripartite complex with Rab3-GTP and synapsin, can ensure that the vesicle cycle is maintained largely by newly endocytosed vesicles by regulating the availability of reserve pool vesicles for release and thus avoiding depletion of the reserve pool.

### Selective tethering of vesicles near the surface membrane enhances the efficiency of the vesicle cycle

Long myosin V tethers are responsible for the initial recruitment and stabilization of vesicles near the presynaptic membrane ^69^ via a direct interaction between vesicle Rab3-GTP and the myosin tail ^70–72^ (Fig. 8 and supplementary video V3). Disruption of the myosin V tethers resulted in a 50% reduction in vesicle usage in experimental studies ^73^. In our model, removing the tethers reduced the rate of vesicle fusion throughout the stimulation period and the total number of fusion events after 45 seconds of 10Hz stimulation by 28% (Fig. 7c). We surmised that the Rab3-myosin V interaction, like the Rab3-RIM interaction, is likely to be inhibited by the binding of tomosyn-1, thus providing a mechanism for the selective tethering of vesicles with free Rab3-GTP (unbound to tomosyn-1) close to the presynaptic membrane. We hypothesized that this ought to improve the efficiency of the vesicle cycle by selectively recruiting recycling vesicles and vesicles released from the cluster with docking propensity, as well as preventing the occlusion of docking sites by vesicles unable to form the crucial Rab3-RIM docking interaction. To test this, we removed the Rab3 dependency in the tethering interaction, such that all vesicles could attach to a tethering path whether or not they contain free (unbound to tomosyn-1) Rab3. Compared to Rab3-dependent selective tethering, non-selective tethering reduced the overall number of vesicles released after 45 seconds of 10 Hz stimulation by 13% whilst simultaneously increasing the number of reserve vesicles used by 48% (Fig. 7a).

**8.**
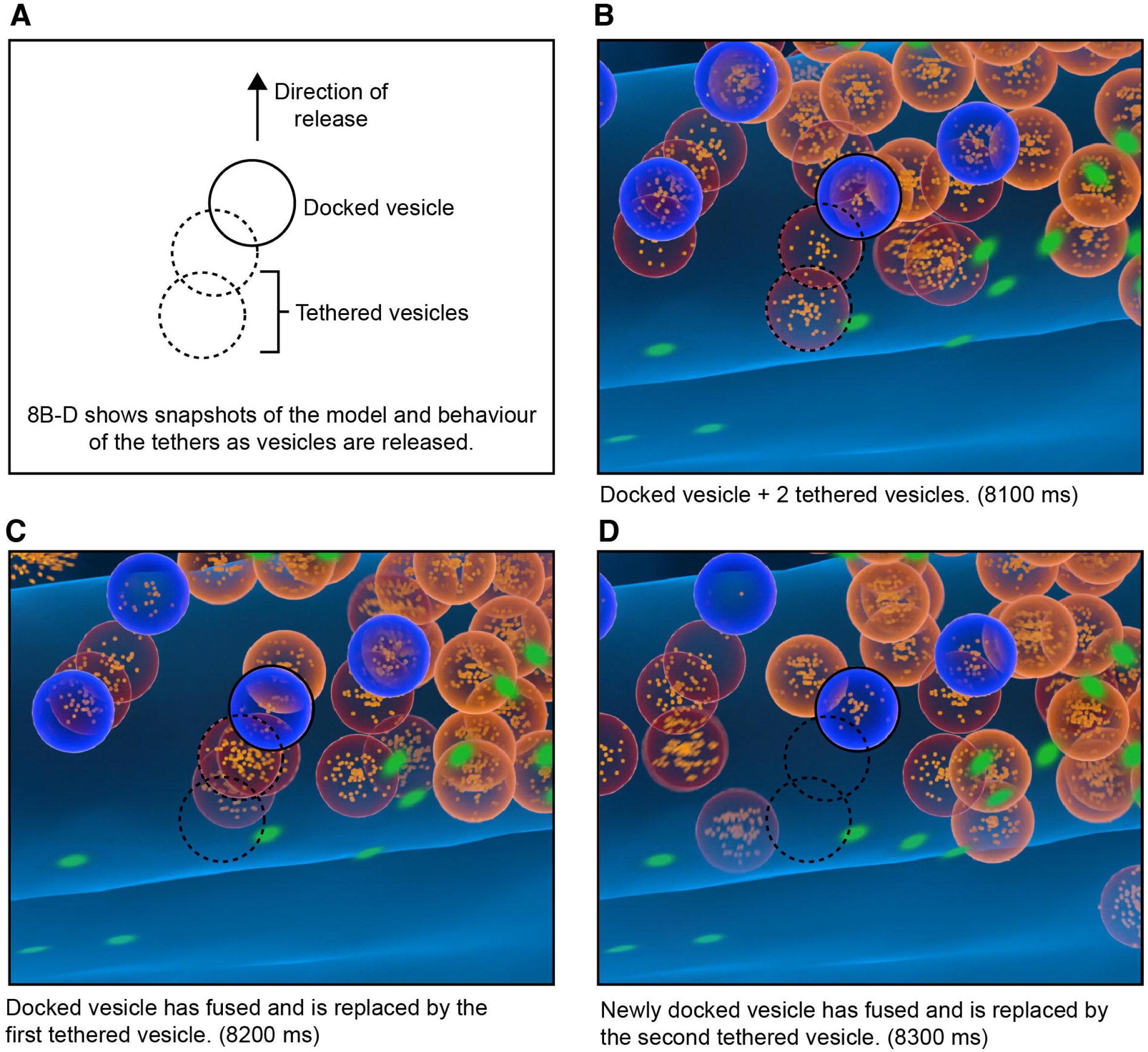
Blender visualization of vesicle tethering. A) Schematic showing tethered vesicles behind a single docked vesicle. B) Snapshot of model showing the tethered vesicles behind a single docked vesicle (see supp. video V3). C) As a docked vesicle fuses, it’s replaced by a tethered vesicle which moves into the vacant docking site. D) The newly docked vesicle fuses and is replaced the next tethered vesicle.

## Methods

### Model Structure

The synaptic bouton morphology was constructed as a tetrahedral mesh using Meshlab and our MultiCompMesher ^74^ derived from an electron micrograph reconstruction of a cultured hippocampal neuron en passant synapse, which also provided the location and area of the active zone ^50^. Cytosolic and vesicle protein copy numbers were derived (where available) from Wilhelm et al 2014 ^18^, with cytosolic and membrane diffusion rates taken from ^50^.

### Simulation of Synaptic Stimulation

All simulations were run on a pre-release version of STEPS 5, which passed all validations presented in Hepburn et al (2023). The hardware used was the ‘Deigo’ cluster at the Okinawa Institute of Science and Technology, on nodes consisting of 2*64 AMD Epyc 7702 @ 2.0GHz cores. The model performance was found to peak at around 256 cores (Hepburn et al 2023), so each individual simulation was run on two full AMD nodes. For 50 seconds model time each simulation took approximately 10 days. Model code will be deposited on the ModelDB site (https://modeldb.science/) upon manuscript publication.

All simulations were run for 5 seconds of model time to reach steady state (to allow vesicle docking and clustering, etc), followed by 45 seconds of stimulation at varying frequencies (5, 10, 20, and 50 Hz). Action potentials were simulated by instantaneous reversal of the membrane potential from –60mV to 40mV for 1ms. Model species, reactions, parameters, and diffusion rates are detailed in Supplementary Tables 1-4.

### Initial Vesicle Populations

All vesicles were populated with vesicle proteins synaptobrevin, synaptotagmin-1, and Rab3-GTP. In addition, to model the buffering of other recycling proteins, vesicles also contain binding sites for complexin, Munc13/18, syndapin, alphaSNAP, and NSF ^75^, with on/off rates tuned to fit the measured proportions of proteins bound to the vesicle cluster ^50^. The only distinction between reserve (initial population: 250) and recycling pool (initial population: 50) vesicles is that the former each contain 10 binding sites for synapsin-1 ^76^ (Fig. 1a). Such that they don’t accumulate synapsin-1 in the timeframe of the simulation ^27^, recycling vesicles (and newly-formed vesicles which become part of the recycling pool ^77^) contain no synapsin-1 binding sites. A single vesicle was placed at each docking site at the beginning of the simulation to allow rapid docking and population of the readily-releasable pool (RRP). The readily-retrievable pool was populated with 40 clathrin-coated pits containing the full complement of vesicle proteins including dynamin ^78^.

### Vesicle Docking

The active zone was populated with eight vesicle docking sites constructed from static RIM1-Munc13 clusters ^79–82^ within the active zone ^83–85^. Calcium channel tethering was modelled by placing four Cav2.1 channels ^58^ in neighboring membrane triangles ^86^.

Vesicle docking is driven by the interaction between vesicle surface Rab3-GTP and the RIM-Munc13 complex. RIM-1 forms a complex with Munc13 which is essential for docking (Fig. 1b and supplementary figure, S1). RIM-1, Munc13, and Rab3-GTP form a tripartite complex which drives docking ^79–82^. Tomosyn-1 binds Rab3-GTP on vesicles, forming a complex that is unable to bind to the RIM1-Munc13 complex, thus preventing docking (Fig. 1c). Tomosyn-1 is phosphorylated by CDK5 and dephosphorylated by calcineurin, with the dephosphorylated form dissociating more rapidly from Rab3-GTP ^31^.

### Vesicle Tethering

Tether paths of length 100nm (allowing 1-2 vesicles to be tethered behind the docked vesicle) were placed at the center along the normal of each docking site ^73, 87^. The requirement for free Rab3-GTP (not complexed with tomosyn-1) on the vesicle for tethering (to model the Rab3-GTP-Myosin V interaction) can be controlled by only allowing vesicles with a specific number of free Rab3-GTP to bind to the tether ^88^. Once attached to a tether path, vesicles will move along the path until reaching the membrane (unless blocked by another docked or tethered vesicle), mimicking retraction of the tether to draw the vesicle into the docking site.

### Vesicle Priming

Munc18 binds tightly to syntaxin-1, locking it into a closed conformation and preventing extraneous binding to SNAP-25 ^89, 90^ (Fig 1b and supplementary figure, S1). Munc13 catalyzes the transfer of the Syntaxin-Munc18 complex into the SNARE complex (binding to SNAP-25) ^89–91^ Kinetics: Zikich 2008). Synaptobrevin (vesicle) is then incorporated into the complex to form the complete SNARE complex ^92, 93^. Vesicle synaptotagmin-1 binds to the SNARE complex, followed by complexin ^2^, fully priming the vesicle for Ca^2+^-triggered fusion.

### Vesicle fusion (exocytosis)

Synaptotagmin-1 comprises two domains (C2A binds three Ca^2+^, and C2B binds two Ca^2+^) ^94, 95^. All five Ca sites must be occupied to trigger conformational change that drives vesicle fusion. Upon Ca-triggered fusion (an exocytosis event in STEPS), any neurotransmitter in the vesicle lumen is expelled into the extracellular space and all vesicle membrane proteins are deposited into the pre-synaptic plasma membrane.

### Post-fusion

Following fusion with the membrane, vesicular proteins disperse (within the bouton only) and then recluster via the adaptor proteins in the pre-assembled pits ^96, 97^ (protein reclustering times fitted to data from Gimber et al 2015) Each vesicle fusion event triggers the generation of a new “pit” in a random position in the peri-endocytic zone for the re-accumulation of dispersed vesicle proteins. The cis-SNARE complex is dismantled by NSF/*α*SNAP ^98, 99^. This is a slow, possibly multi-step, reaction taking ∼6 seconds per SNARE complex ^100^.

### Vesicle retrieval and recycling

Newly-formed pits contain adaptors for synaptotagmin-1 (AP2) and synaptobrevin (AP180), which are responsible for the reclustering and accumulation of these vesicle proteins in the readily-retrievable pool following vesicle fusion and protein dispersion. As well as triggering vesicle fusion, Ca^2+^ also activates calcineurin (via calmodulin). Active calcineurin dephosphorylates (and thus activates) dynamin. Active dynamin is responsible for the last step in endocytosis of vesicles from the readily-retrievable pool ^15, 60^. Only pits with their full complement of synaptotagmin-1, synaptobrevin, and Rab3-GTP can be retrieved from the readily-retrievable pool. Once formed, the newly-retrieved vesicles become part of the recycling pool and can dock and fuse at the active zone.

### Vesicle Clustering Model

Synapsins accumulate on the vesicle surface ^101, 102^ and synapsin-synapsin interactions cause vesicles to cluster ^29^. Vesicle clustering is not dependent on synapsin-actin interactions ^68^, but static actin proteins in the cytosol act as seeds for the vesicle cluster. Synapsin-1 is phosphorylated by PKA and dephosphorylated by PP2A (Fig. 1c). The phosphorylated form of synapsin-1 dissociates more rapidly from both the vesicles and into the monomeric form ^96, 103, 104^. Vesicle-bound synapsin-1 also forms a stable heterotrimer with the tomosyn-Rab3-GTP complex ^31^.

## Discussion

Our detailed spatial model of the complete vesicle cycle reveals how a complex interplay of vesicle and cytosolic proteins, molecular tethering at the plasma membrane, and the maintenance of a dynamic vesicle cluster, generate a remarkably robust recycling system, able to maintain vesicle release at frequencies well beyond what could be considered physiological for hippocampal neurons. In particular, we reveal how Rab3, synapsin-1, and tomosyn-1 work together to maintain and regulate the release of vesicles from the vesicle cluster and their propensity for release, thus providing a mechanism for the dynamic segregation of the recycling and reserve pools depending on need. Furthermore, we show how selective tethering of vesicles at the membrane can improve the efficiency of the vesicle cycle by preventing the docking of vesicles with a low propensity for release and reducing the number of vesicles recruited from the reserve pool. Also, despite its limited availability, saturation of dynamin – controlling the final step in endocytosis – appears unlikely to be an issue even at non-physiological firing frequencies for up to a minute.

At all frequencies studied, endocytosis followed exocytosis closely and, following depletion of the recycling pool, the majority of fusion events were from recycled vesicles, indicating that vesicle recycling is mainly responsible for maintaining synaptic transmission over time. This dovetails with experimental studies demonstrating that, at relatively low frequencies, recycling can be maintained using only a small proportion of the entire vesicle pool ^11, 13, 105^, as well as a study showing that newly-endocytosed vesicles are preferentially utilized over more mature vesicles ^28^. However, as the firing rate and stimulation time increases, the recycling pathway becomes overwhelmed and is unable to compensate for the rapid rate of vesicle fusion: the recycling pool is depleted and vesicle material begins to accumulate in the membrane. In the absence of a reserve pool, this would lead to a gradual decrease in release probability, threatening the fidelity of synaptic transmission. However, dispersion of the vesicle cluster, together with CaN-mediated dephosphorylation of tomosyn-1 and destabilization of the Rab3-tomosyn-synapsin tripartite complex, releases vesicles from the reserve pool for docking and fusion. Once a reserve vesicle is utilized and recycled, it becomes part of – and thus increases the size of – the recycling pool, further increasing the availability of vesicles for release without the requirement for direct coupling of exocytosis and endocytosis. The purpose of the reserve pool is not simply to supply additional vesicles when the recycling pool is depleted, but to release vesicles into the vesicle cycle and increase the number of actively recycling vesicles (following fusion and retrieval), which allows the cycle to maintain a higher rate of fusion and retrieval over time.

Although considering vesicles as being part of separate recycling and reserve pools is an instructive simplification, the distinction between the recycling and reserve vesicle pools doesn’t appear to be sharply delineated ^9^. In hippocampal neurons, all vesicles, including those of the putative reserve pool, will eventually fuse during mild stimulation over extended periods of time ^9, 17^. This suggests that the likelihood of a particular vesicle being utilized is largely probabilistic, with the recycling pool simply comprising those vesicles with a relatively high propensity for docking and release, although this propensity can change as a vesicle matures or, as we have shown, dependent on synaptic activity and the pattern of vesicle protein phosphorylation ^28^.

Whilst it’s ostensibly reasonable to suppose that release of vesicles from the cluster is alone responsible for rendering them available for docking and release, effectively transferring them from the reserve to the recycling pool, the vesicle cluster often lies close to the active zone and both types of vesicles are intermixed. As such, clustering alone is unlikely to provide control over pool segregation and recruitment. However, synapsin-1 and tomosyn-1 can work together to both maintain the population of reserve vesicles within the synaptic bouton and to control their propensity for fusion and thus, following retrieval, release into the recycling pool. By forming a stable tripartite complex, Rab3, tomosyn-1, and synapsin-1 can lock vesicles in a non-fusion-competent state within the vesicle cluster. PKA-mediated phosphorylation of synapsin causes dispersal of the vesicle cluster and promotes loss of tomosyn-1 from Rab3, effectively releasing vesicles from the reserve pool. At high firing rates, this is further promoted by CaN-dependent dephosphorylation of tomosyn-1, which disrupts the tomosyn-1-synapsin interaction. Although our model shows that tomosyn-1 doesn’t appear to regulate the overall availability of vesicles for release, it is able to control the proportion of vesicles from the reserve pool that are utilized. Blocking PKA-mediated dispersion of the cluster or increasing tomosyn-1 copy number prevents reserve vesicle recruitment and compromises vesicle release at higher firing rates.

In addition to efficient recycling of vesicle material following exocytosis, stable vesicle release also depends upon the rapid replenishment of vacant docking sites. Whilst recycling vesicles are observed to diffuse freely through the cytosol, intermixing with the vesicle cluster, relying on random diffusion alone to bring vesicles close to the active zone docking sites is inefficient in replenishing fused vesicles. Long myosin V tethers are responsible for initial recruitment and stabilization of vesicles much further from the membrane ^69^, ensuring the RRP can be replenished in a timely manner during repetitive stimulation, with myosin V disruption resulting in a 50% reduction in vesicle usage in experimental studies ^73^ and by 28% in our model. Since both the recycling and reserve pool vesicles are packed into the bouton close to the active zone ^14, 23^, it’s likely that tethering should be somewhat selective for recycling pool vesicles to avoid clogging the active zone with reserve vesicles that aren’t viable for release. And, since Rab3-GTP interacts with myosin V (potentially via an unidentified effector protein)^71, 88^, it’s plausible that this long-distance tethering is also dependent on free Rab3-GTP. Once a vesicle is moved to the docking site, Rab3-myosin V interactions can be displaced by the Rab3-RIM1-Munc13 docking interactions. In support of this model, selective tethering both enhances the rate of vesicle release over time whilst also reducing the number of vesicles recruited from the reserve pool to maintain synaptic release. Reserve pool vesicle recruitment is not cost-free, since released vesicles must ultimately be returned to the reserve pool cluster by the slow accumulation of synapsin to avoid being lost from the bouton. Selective tethering allows a smaller number of recycling vesicles to maintain the vesicle cycle with minimal reserve pool recruitment at physiological firing rates.

Whilst our model captures all phases of the vesicle cycle, it is by no means complete. For example, we don’t take into account possible involvement of “superprimed” vesicles ^106^, nor tomosyn-1’s role in inhibiting vesicle priming by antagonizing Munc13 and inhibiting SNARE complex formation ^107–111^. However, our unique vesicle modeling technology will provide rich opportunities for studying these and other mechanisms involved in the highly complex and dynamic regulation of the synaptic vesicle cycle.

## Acknowledgements.

We are extremely grateful to Pavel Puchenkov of the Scientific Data Visualization Support service, Research Support Division, Okinawa Institute of Science and Technology Graduate University for invaluable help in designing and coding the visualization of the model in Blender.

This work was funded in part by a Grants-in-Aid for Scientific Research (KAKENHI) grant awarded to ARG, IH, and EDS, and in part by the Okinawa Institute of Science and Technology Graduate University.

## Supplementary Figure and Video Legends

S1. A-F shows the docking and priming steps modelled, from initial docking via the Rab3-GTP to RIM-Muncs13 interaction through to the fully primed vesicle ready for calcium-triggered fusion.

S2. Distribution of vesicle recycling times (time from fusion until endocytosis) following 500 vesicle fusion events. Mean recycling time = 6.8 seconds.

S3. The effect of blocking calcineurin (CaN)-mediated dephosphorylation of Tomosyn-1 on release of vesicles from reserve pool.

V1. A toy cylindrical model showing the dynamic clustering of 250 synaptic vesicles. Clustered vesicles are blue, freely diffusing vesicles are shown in red.

V2. Visualization showing dispersal of the vesicle cluster over ∼30 seconds during repetitive firing. Note that vesicle fusion is blocked and action potentials are not visualized for clarity.

V3. Visualization of the complete vesicle model, showing the dynamic vesicle cluster, vesicle fusion and endocytosis, and the replenishment of fused vesicles. Each white “flash” indicates an action potential. See Fig. 2 of the main text for key to vesicle types and Fig. 8 for details of tethering and vesicle replenishment. Note that the release of neurotransmitter following vesicle fusion is represented using a Blender particle system, is for visualization purposes only, and does not represent model data.

