## Supplementary figures and images for "Dynamic Regulation of Vesicle Pools in a Detailed Spatial Model of the Complete Synaptic Vesicle Cycle"

### Supp. Fig. 1

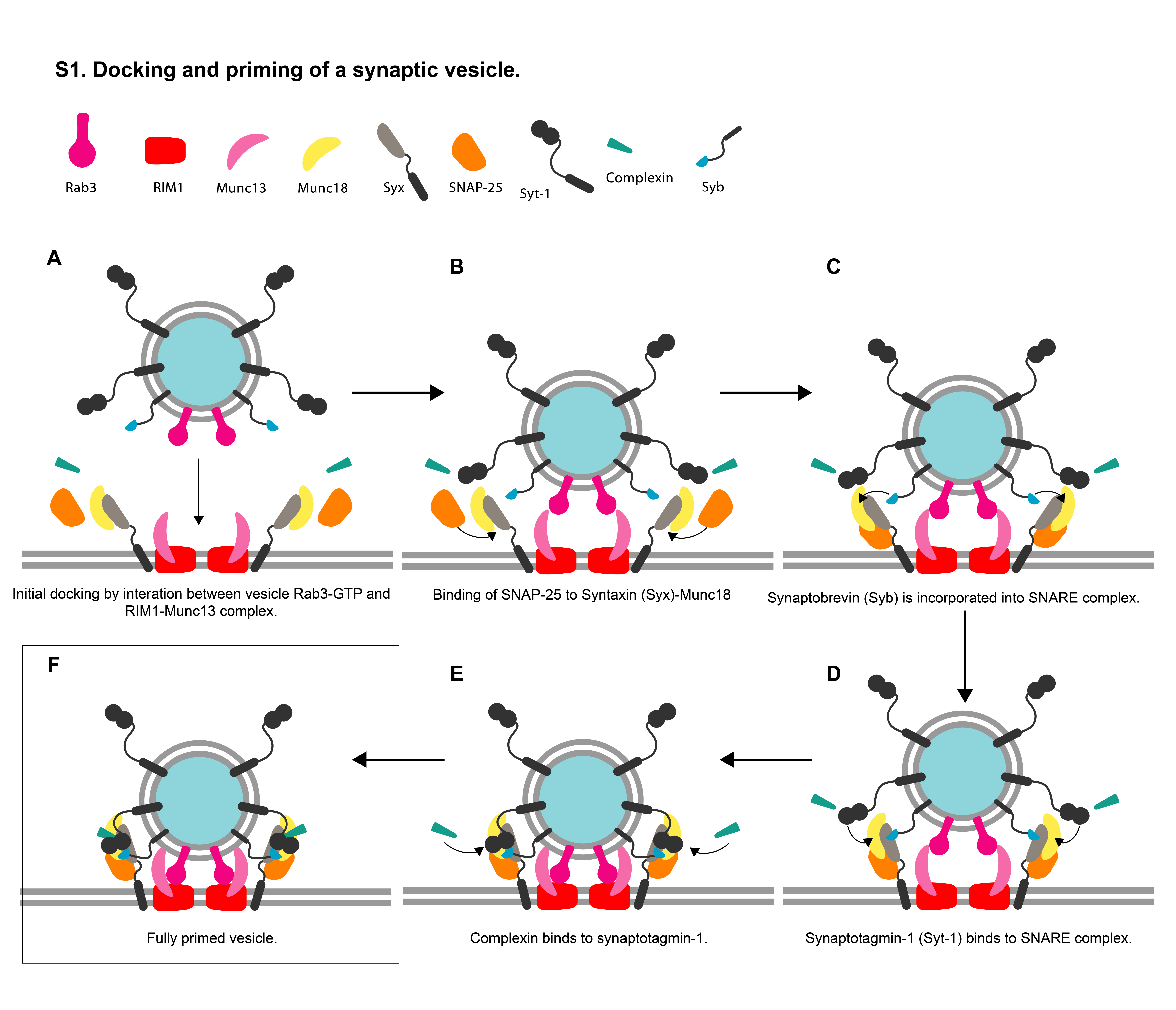

### Supp. Fig. 2

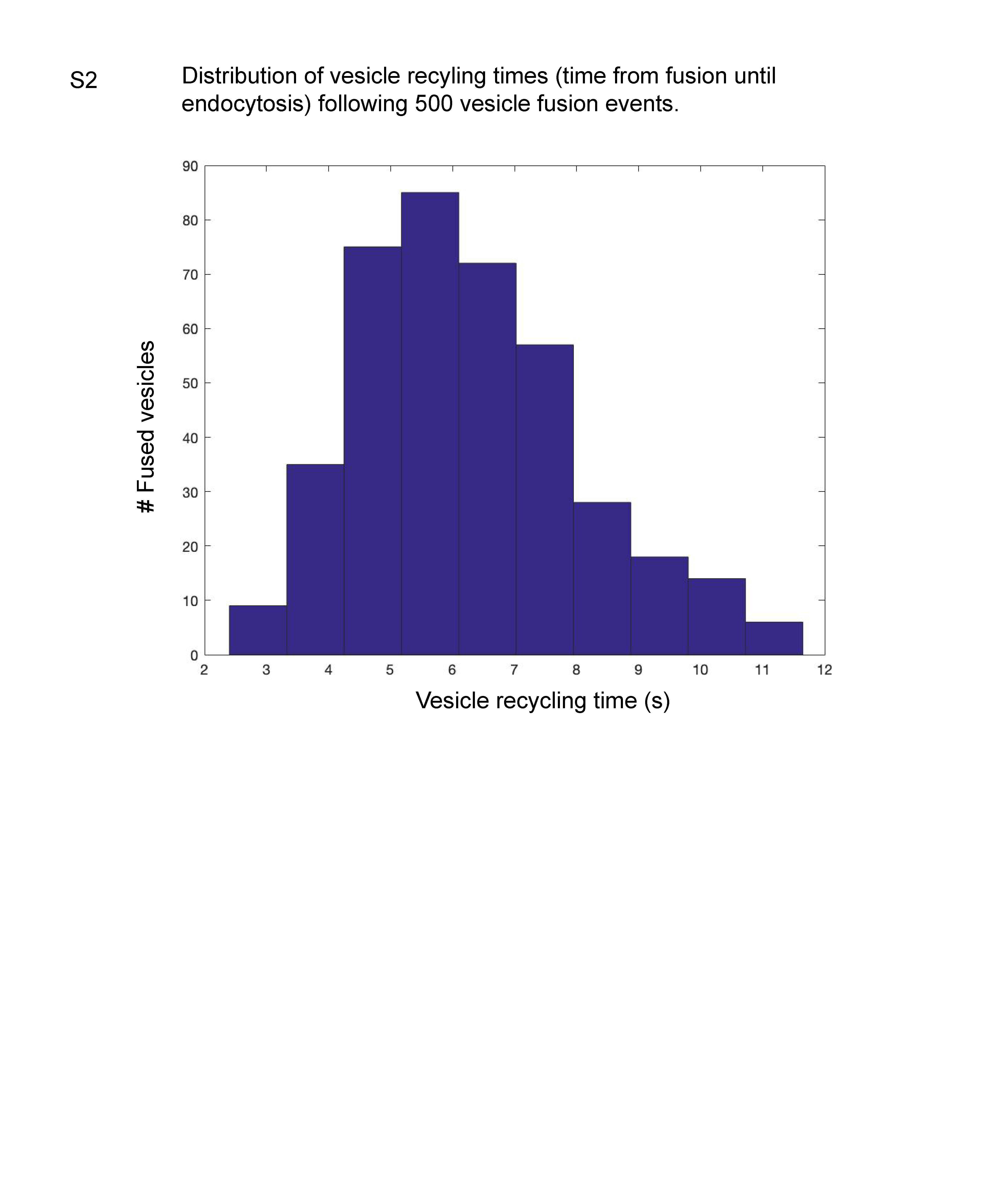

### Supp. Fig. 3

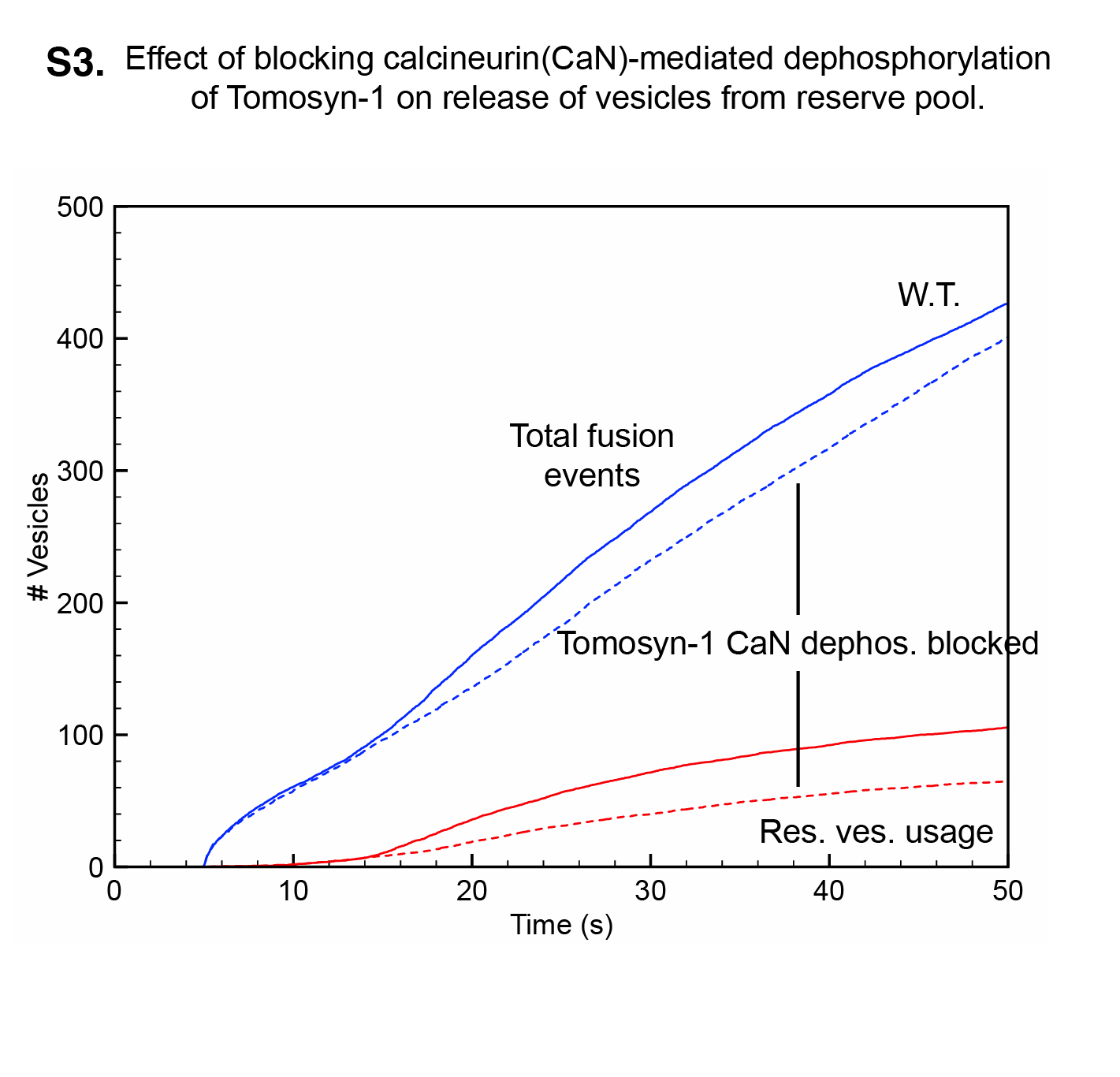
