## Supplementary material for "Dynamic Regulation of Vesicle Pools in a Detailed Spatial Model of the Complete Synaptic Vesicle Cycle": Supp. Tables 1-4

### Model Species, Parameters, and Reactions

Supplementary Table 1

| Model Species | Full name | Initial concentration (µM)/(number) | Ref. |
|---|---|---|---|
| Ca (cytosol) | Calcium | 0.045 | Doi 2005; Antunes 2012; Bartol 2015 |
| CBhi', | Calbindin (high affinity) | 0.99 | Doi 2005; Antunes 2012; Bartol 2015 |
| CBlo | Calbindin (low affinity) | 0.99 | Doi 2005; Antunes 2012; Bartol 2015 |
| CRTT | | 0.1976 | Faas 2007 |
| CRind | | 0.494 | Faas 2007 |
| PV | Parvalbumin | 4.55 | Lee 2000 |
| CaPV | | 8.4 | Antunes 2012 |
| MgPV | | 30.45 | Antunes 2012 |
| AC18 | Adenylyl cyclase | 0.2 | Bhalla 1999 |
| R2C2 | PKA (inactive form) | 0.5 | Bhalla 1999 |
| SERCA | See code | SERCA_count | Doi 2005; Antunes 2012; Bartol 2015 |
| PMCA_P0 | See code | PMCA_count | Bartol 2015 |
| Ca (ER) | Calcium | 150 | Doi 2005 |
| Rab3 | Rab3-GTP | 10 per vesicle | Takamori 2006 |
| SYB | Synaptobrevin | 69 per vesicle | Takamori 2006 |
| syt | Synaptotagmin-1 | 27 per vesicle | Takamori 2006 |
| DYNp | Dynamin | 1 per raft in RRetP | Ross 2011; Chugh 2006; Jimah_2019 |
| RIM_M13 | RIM1_Munc13 complex | 4 per docking site | Imig 2014, Murthy 1999 |
| CaP_m0 | P-type Ca channel | 4 per docking site | Althof 2015 |
| M18 | Munc18 | 28.4 | Wilhelm 2014 |
| M13 | Munc13 | 10.36 | Wilhelm 2014 |
| CXN | Complexin | 16.59 | Wilhelm 2014 |
| aSNAP | alphaSNAP | 7.68 | Wilhelm 2014 |
| NSF | NSF | 27.14 | Wilhelm 2014 |
| Synapsin | Synapsin | 156 | Wilhelm 2014 |
| PP2A | PP2A | 0.5 | Bhalla 1999; Gallimore 2018 |
| SYX | Syntaxin | (32153) | Wilhelm 2014 |
| SNP25 | SNAP25 | (42697) | Wilhelm 2014 |

| | | | |
|---|---|---|---|
| CaM_N_0_C_0 | Calmodulin (free form) | 60 | Kawaguchi 2013; Kitagawa 2009; Wilhelm 2014 |
| CaN | Calcineurin | 5 | Kitagawa 2009 |
| CDK5 | CDK5 | 1 | Shin 2019 |
| Rab3 | Rab3-GTP | 125.8 | Wilhelm 2014 |
| SYN1 | Syndapin-1 | 21.37 | Wilhelm 2014 |
| TOMO_p | Tomosyn-1 (phospho form) | 1500 | Barak 2010; Fujita 1998 |

Supplementary Table 2

| Species | Cytosolic diffusion constant (μm2/s) | Membrane diffusion constant (μm2/s) |
|---|---|---|
| **Unlisted species don't diffuse.** | **Obtained from Reshetniak 2020 (where available)** | **Obtained from Reshetniak 2020 (where available)** |
| Vesicles | 0.01 | |
| Ca (cytosol) | 223 | |
| CBhi', | 2.0 | |
| CBlo | 2.0 | |
| CRTT | 2.0 | |
| CRind | 2.0 | |
| PV | 2.0 | |
| CaPV | 2.0 | |
| MgPV | 2.0 | |
| AC18 | 2.0 | |
| R2C2 | 2.0 | |
| SERCA | | 0.25 |
| PMCA_P0 | | 0.25 |
| Rab3 | 1.7793 | 0.25 |
| SYB | | 0.3245 |
| syt | | 0.25 |
| M18 | 2.2357 | |
| M13 | 2.2796 | |
| CXN | 1.7793 | |
| aSNAP | 2.1959 | |
| NSF | 2.2788 | |
| Synapsin | 2.65 | |
| PP2A | 2.0 | |
| SYX | 0.2429 | |
| SNP25 | | 0.651 |
| CaM_N_0_C_0 | 0.25 | |
| CaN | 0.25 | |
| CDK5 | 0.25 | |
| SYN1 | 2.1959 | |
| TOMO_p | 1.7793 | |

Supplementary Table 3

| Parameter | Value | Units | Notes and ref. |
|---|---|---|---|
| kon_SNARE_syt_CXN_ca1 | 1000 | $\mu M^{-1}s^{-1}$ | Millet 2002; Davis 1999; Hui 2005 |
| koff_SNARE_syt_CXN_ca1 | 1270 | $s^{-1}$ | " " |
| kon_SNARE_syt_CXN_ca2 | 1000 | $\mu M^{-1}s^{-1}$ | " " |
| koff_SNARE_syt_CXN_ca2 | 227670 | $s^{-1}$ | " " |
| kon_SNARE_syt_CXN_ca3 | 1000 | $\mu M^{-1}s^{-1}$ | " " |
| koff_SNARE_syt_CXN_ca3 | 12370 | $s^{-1}$ | " " |
| kon_SNARE_syt_CXN_bca1 | 1000 | $\mu M^{-1}s^{-1}$ | " " |
| koff_SNARE_syt_CXN_bca1 | 50000 | $s^{-1}$ | " " |
| kon_SNARE_syt_CXN_bca2 | 1000 | $\mu M^{-1}s^{-1}$ | " " |
| koff_SNARE_syt_CXN_bca2 | 25780 | $s^{-1}$ | " " |
| koff_SNARE_syt_CXN_ca1_s | 1000 | $s^{-1}$ | " " |
| koff_SNARE_syt_CXN_ca2_s | 2000 | $s^{-1}$ | " " |
| koff_SNARE_syt_CXN_ca3_s | 5000 | $s^{-1}$ | " " |
| kon_rim_rab3 | 1000 | $\mu M^{-1}s^{-1}$ | Model calibration |
| koff_rim_rab3 | 0 | $s^{-1}$ | Docking assumed to be irreversible once vesicle is positioned at the dock site. |
| kon_syx_m18 | 5 | $\mu M^{-1}s^{-1}$ | Burkhardt 2008 |
| koff_syx_m18 | 0.0011 | $s^{-1}$ | " " |
| kon_m13_syx | 20 | $\mu M^{-1}s^{-1}$ | Mezer 2004 |
| koff_m13_syx | 2.6 | $s^{-1}$ | " " |
| kon_syx_snp25 | 10 | $\mu M^{-1}s^{-1}$ | Zikich 2008; Coppola_2001 |
| koff_syx_snp25 | 1.26 | $s^{-1}$ | " " |
| kon_syb_syx | 2.35 | $\mu M^{-1}s^{-1}$ | Lai 2017; Shu 2020; Walter 2010 |
| koff_syb_syx | 0.0047 | $s^{-1}$ | " " |
| kon_syt_snare | 57 | $\mu M^{-1}s^{-1}$ | Mezer 2004 |
| koff_syt_snare | 0.24 | $s^{-1}$ | " " |
| kon_cxn_snare | 30 | $\mu M^{-1}s^{-1}$ | Mezer 2004; Li 2007; Bowen 2005 |
| koff_cxn_snare | 0.33 | $s^{-1}$ | " " |
| kon_snare_snap | 0.17 | $\mu M^{-1}s^{-1}$ | Rickman 2002 |
| koff_snare_snap | 0.26 | $s^{-1}$ | " " |
| kon_nsf_snap | 20 | $\mu M^{-1}s^{-1}$ | Vivona 2013; Cipriano 2013 |

| | | | |
|---|---|---|---|
| kcat_nsf | 0.116 | $s^{-1}$ | Vivona 2013; Cipriano 2013 |
| kon_cam_NT | 770 | $\mu M^{-1}s^{-1}$ | Faas 2011; Sun 2010 |
| kon_cam_CT | 84 | $\mu M^{-1}s^{-1}$ | " " |
| kon_cam_NR | 32000 | $\mu M^{-1}s^{-1}$ | " " |
| kon_cam_CR | 25 | $\mu M^{-1}s^{-1}$ | " " |
| koff_cam_NT | $1.6x10^5$ | $s^{-1}$ | " " |
| koff_cam_CT | $2.2x10^4$ | $s^{-1}$ | " " |
| koff_cam_CR | 2600 | $s^{-1}$ | " " |
| koff_cam_NR | 6.5 | $s^{-1}$ | " " |
| kon_can_cam | 46 | $\mu M^{-1}s^{-1}$ | Quintana 2005 |
| koff_can_cam | 1.2 | $s^{-1}$ | " " |
| kon_can_dyn | 10 | $\mu M^{-1}s^{-1}$ | Model calibration |
| koff_can_dyn | 1 | $s^{-1}$ | " " |
| kcat_can | 0.5 | $s^{-1}$ | " " |
| kon_dyn_syn | 100 | $\mu M^{-1}s^{-1}$ | Smillie 2005 |
| koff_dyn_syn | 1 | $s^{-1}$ | Anggono 2006 |
| kon_cdk_dyn | 3 | $\mu M^{-1}s^{-1}$ | Liu 2010 |
| koff_cdk_dyn | 10 | $s^{-1}$ | " " |
| kcat_cdk | 0.5 | $s^{-1}$ | " " |
| kon_syb_ap180 | 2000 | $\mu M^{-1}s^{-1}$ | Assumed to be fast and irreversible. Gordon 2014 |
| kon_syb_ap2 | 2000 | $\mu M^{-1}s^{-1}$ | Diril 2006 |
| kon_ac_cam | 500 | $\mu M^{-1}s^{-1}$ | Wang 2003 |
| koff_ac_cam | 0.1 | $s^{-1}$ | " " |
| kcat_ac18 | 18 | $s^{-1}$ | Wu 1993; Bhalla 1999 |
| kdeg_camp | 1 | $s^{-1}$ | This paper. |
| kon_r2c2_camp1 | 2 | $\mu M^{-1}s^{-1}$ | Buxbaum 1989; Bhalla 1999 |
| koff_r2c2_camp1 | 0.75 | $s^{-1}$ | " " |
| kon_r2c2_camp2bb | 1 | $\mu M^{-1}s^{-1}$ | " " |
| koff_r2c2_camp2bb | 1.5 | $s^{-1}$ | " " |
| kon_r2c2_camp2ab | 10 | $\mu M^{-1}s^{-1}$ | " " |
| koff_r2c2_camp2ab | 7.5 | $s^{-1}$ | " " |
| kon_r2c2_camp3bb | 20 | $\mu M^{-1}s^{-1}$ | " " |
| koff_r2c2_camp3bb | 7.5 | $s^{-1}$ | " " |
| kon_r2c2_camp3ab | 1 | $\mu M^{-1}s^{-1}$ | " " |
| koff_r2c2_camp3ab | 0.75 | $s^{-1}$ | " " |
| kon_r2c2_camp4 | 10 | $\mu M^{-1}s^{-1}$ | " " |
| koff_r2c2_camp4 | 15 | $s^{-1}$ | " " |
| kact_pka_23 | 0.005 | $s^{-1}$ | " " |

| | | | |
|---|---|---|---|
| kdeact_pka_23 | $5 \times 10^6$ | $s^{-1}$ | " " |
| kact_pka_4 | 6 | $s^{-1}$ | " " |
| kdeact_pka_4 | $5 \times 10^6$ | $s^{-1}$ | " " |
| kon_r2c_camp3 | 1 | $\mu M^{-1}s^{-1}$ | " " |
| koff_r2c_camp3 | 0.75 | $s^{-1}$ | " " |
| kon_r2c_camp4 | 10 | $\mu M^{-1}s^{-1}$ | " " |
| koff_r2c_camp4 | 7.5 | $s^{-1}$ | " " |
| kact_pka_r2c | 3 | $s^{-1}$ | " " |
| kdeact_pka_r2c | $10 \times 10^6$ | $s^{-1}$ | " " |
| kon_PKA_syn | 10 | $\mu M^{-1}s^{-1}$ | Model calibration |
| koff_PKA_syn | 1 | $s^{-1}$ | " " |
| kcat_PKA_syn | 10 | $s^{-1}$ | " " |
| kon_pp2a_synapsin | 2 | $\mu M^{-1}s^{-1}$ | Gallimore 2018 |
| koff_pp2a_synapsin | 8 | $s^{-1}$ | " " |
| kcat_pp2a_syn | 2 | $s^{-1}$ | " " |
| kon_synapsin_actin | 1000 | $\mu M^{-1}s^{-1}$ | Model calibration |
| koff_synapsin_actin | 1 | $s^{-1}$ | " " |
| koff_synapsin_actin_p | 1000 | $s^{-1}$ | " " |
| kon_cdk_tomo | 3 | $\mu M^{-1}s^{-1}$ | " " |
| koff_cdk_tomo | 10 | $s^{-1}$ | " " |
| kcat_cdk_tomo | 0.5 | $s^{-1}$ | " " |
| kon_can_tomo | 1 | $\mu M^{-1}s^{-1}$ | " " |
| koff_can_tomo | 10 | $s^{-1}$ | " " |
| kcat_can_tomo | 1 | $s^{-1}$ | " " |
| kon_rab3_TOMO_p | 1 | $\mu M^{-1}s^{-1}$ | " " |
| koff_rab3_TOMO_p | 10 | $s^{-1}$ | " " |
| kon_TOMO_p_synapsin | 100 | $\mu M^{-1}s^{-1}$ | " " |
| koff_TOMO_p_synapsin | 1 | $s^{-1}$ | " " |
| kon_TOMO_synapsin | 100 | $\mu M^{-1}s^{-1}$ | " " |
| koff_TOMO_synapsin | 100 | $s^{-1}$ | Increased off-rate with dephos. |
| kon_synapsin_dimer | 10 | $\mu M^{-1}s^{-1}$ | This paper. |
| koff_synapsin_dimer | 10 | $s^{-1}$ | " " |
| kon_synapsin_dimer_p | 10000 | $\mu M^{-1}s^{-1}$ | Increased off-rate with phos. |
| kon_synapsin_ves | 10.7 | $\mu M^{-1}s^{-1}$ | Calibration against Reshetniak 2020 |
| koff_synapsin_ves | 0.123 | $s^{-1}$ | " " |
| koff_synapsin_p_ves | 123 | $s^{-1}$ | Increased off-rate with phos. |
| kon_cxn_ves | 1.2 | $\mu M^{-1}s^{-1}$ | Zdanowicz 2017 |
| koff_cxn_ves | 1000 | $s^{-1}$ | " " |

| | | | |
|---|---|---|---|
| kon_rab3_ves | 10 | $\mu M^{-1}s^{-1}$ | Calibration against Reshetniak 2020 |
| koff_rab3_ves | 1 | $s^{-1}$ | " " |
| kon_m13_ves | 1.6 | $\mu M^{-1}s^{-1}$ | " " |
| koff_m13_ves | 1000 | $s^{-1}$ | " " |
| kon_m18_ves | 1.6 | $\mu M^{-1}s^{-1}$ | " " |
| koff_m18_ves | 1000 | $s^{-1}$ | " " |
| kon_syn1_ves | 2.8 | $\mu M^{-1}s^{-1}$ | " " |
| koff_syn1_ves | 1000 | $s^{-1}$ | " " |
| kon_asnap_ves | 2.8 | $\mu M^{-1}s^{-1}$ | " " |
| koff_asnap_ves | 1000 | $s^{-1}$ | " " |
| kon_nsf_ves | 0.9 | $\mu M^{-1}s^{-1}$ | " " |
| koff_nsf_ves | 1000 | $s^{-1}$ | " " |

Supplementary Table 4

| Reaction | Parameter |
|---|---|
| **Synaptotagmin (Syt) model** | |
| Syt + Ca ↔ Syt_Ca | kon_SNARE_syt_CXN_ca1<br>koff_SNARE_syt_CXN_ca1 |
| Syt_Ca + Ca ↔ Syt_Ca2 | kon_SNARE_syt_CXN_ca2<br>koff_SNARE_syt_CXN_ca2 |
| Syt_Ca2 + Ca ↔ Syt_Ca3 | kon_SNARE_syt_CXN_ca3<br>koff_SNARE_syt_CXN_ca3 |
| Syt + Ca ↔ Syt_bCa | kon_SNARE_syt_CXN_bca1<br>koff_SNARE_syt_CXN_bca1 |
| Syt_bCa + Ca ↔ Syt_bCa2 | kon_SNARE_syt_CXN_bca2<br>koff_SNARE_syt_CXN_bca2 |
| Syt_bCa + Ca ↔ Syt_Ca_bCa | kon_SNARE_syt_CXN_ca1<br>koff_SNARE_syt_CXN_ca1 |
| Syt_Ca_bCa + Ca ↔ Syt_Ca2_bCa | kon_SNARE_syt_CXN_ca2<br>koff_SNARE_syt_CXN_ca2 |
| Syt_Ca2_bCa + Ca ↔ Syt_Ca3_bCa | kon_SNARE_syt_CXN_ca3<br>koff_SNARE_syt_CXN_ca3 |
| Syt_Ca + Ca ↔ Syt_Ca_bCa | kon_SNARE_syt_CXN_bca1<br>koff_SNARE_syt_CXN_bca1 |
| Syt_Ca_bCa + Ca ↔ Syt_Ca_bCa2 | kon_SNARE_syt_CXN_bca2<br>koff_SNARE_syt_CXN_bca2 |
| Syt_Ca2 + Ca ↔ Syt_Ca2_bCa | kon_SNARE_syt_CXN_bca1<br>koff_SNARE_syt_CXN_bca1 |
| Syt_Ca2_bCa + Ca ↔ Syt_Ca2_bCa2 | kon_SNARE_syt_CXN_bca2<br>koff_SNARE_syt_CXN_bca2 |
| Syt_Ca3 + Ca ↔ Syt_Ca3_bCa | kon_SNARE_syt_CXN_bca1<br>koff_SNARE_syt_CXN_bca1 |
| Syt_Ca3_bCa + Ca ↔ Syt_Ca3_bCa2 | kon_SNARE_syt_CXN_bca2<br>koff_SNARE_syt_CXN_bca2 |

| | |
|---|---|
| Syt_bCa2 + Ca ↔ Syt_Ca_bCa2 | kon_SNARE_syt_CXN_ca1<br>koff_SNARE_syt_CXN_ca1_s |
| Syt_Ca_bCa2 + Ca ↔ Syt_Ca2_bCa2 | kon_SNARE_syt_CXN_ca2<br>koff_SNARE_syt_CXN_ca2_s |
| Syt_Ca2_bCa2 + Ca ↔ Syt_Ca3_bCa2 | kon_SNARE_syt_CXN_ca3<br>koff_SNARE_syt_CXN_ca3_s |
| **Vesicle Docking.** | |
| Rab3(vesicle) + RIM_M13(membrane) ↔ RIM_M13_Rab3 | kon_rim_rab3<br>koff_rim_rab3 |
| **Vesicle Priming I** | |
| SYX + M18 ↔ SYX_M18 | kon_syx_m18<br>koff_syx_m18 |
| RIM_M13_Rab3 + SYX_M18 ↔ RIM_M13_Rab3_SYX_M18 | kon_m13_syx<br>koff_m13_syx |
| RIM_M13_Rab3_SYX_M18 SNP25 ↔ RIM_M13_Rab3_SYX_M18_SNP25 | kon_syx_snp25<br>koff_syx_snp25 |
| RIM_M13_Rab3_SYX_M18_SNP25 + SYB ↔ SNARE | kon_syb_syx<br>koff_syb_syx |
| **Vesicle Priming II** | |
| SNARE + Syt ↔ SNARE_Syt | kon_syt_snare<br>koff_syt_snare |
| SNARE_Syt + CXN ↔ SNARE_syt_CXN | kon_cxn_snare<br>koff_cxn_snare |
| **Vesicle Fusion (exocytosis)** | |
| Requires 2xSNARE_syt_CXN_Ca3_bCa2 | 3e03 |
| Requires 3xSNARE_syt_CXN_Ca3_bCa2 | 3e04 |
| Requires 4xSNARE_syt_CXN_Ca3_bCa2 | 3e05 |
| **NSF dismantling of SNARE complex** | |
| cisSNARE + aSNAP ↔ SNARE_aSNAP | kon_snare_snap<br>koff_snare_snap |
| SNARE_aSNAP + NSF → SNARE_aSNAP_NSF | kon_nsf_snap |
| SNARE_aSNAP_NSF → SYB + SYX + SNP25 + NSF + aSNAP + M18 | kcat_nsf |
| **Ca_Calmodulin Model** | |
| **Ca ON Reactions** | |
| CaM_N_0_C_0 + Ca → CaM_N_1_C_0 | 2*kon_cam_NT |
| CaM_N_0_C_0 + Ca → CaM_N_0_C_1 | 2*kon_cam_CT |
| CaM_N_1_C_0 + Ca → CaM_N_1_C_1 | 2*kon_cam_CT |
| CaM_N_1_C_0 + Ca → CaM_N_2_C_0 | kon_cam_NR |
| CaM_N_0_C_1 + Ca → CaM_N_0_C_2 | kon_cam_CR |
| CaM_N_0_C_1 + Ca → CaM_N_1_C_1 | 2*kon_cam_NT |
| CaM_N_1_C_1 + Ca → CaM_N_2_C_1 | kon_cam_NR |
| CaM_N_1_C_1 + Ca → CaM_N_1_C_2 | kon_cam_CR |
| CaM_N_2_C_0 + Ca → CaM_N_2_C_1 | 2*kon_cam_CT |
| CaM_N_0_C_2 + Ca → CaM_N_1_C_2 | 2*kon_cam_NT |
| CaM_N_1_C_1 + Ca → CaM_N_2_C_1 | kon_cam_NR |

| | |
|---|---|
| CaM_N_1_C_1 + Ca → CaM_N_1_C_2 | kon_cam_CR |
| CaM_N_2_C_1 + Ca → CaM_N_2_C_2 | kon_cam_CR |
| CaM_N_1_C_2 + Ca → CaM_N_2_C_2 | kon_cam_NR |
| **Ca OFF Reactions** | |
| CaM_N_1_C_0 → CaM_N_0_C_0 + Ca | koff_cam_NT |
| CaM_N_0_C_1 → CaM_N_0_C_0 + Ca | koff_cam_CT |
| CaM_N_1_C_1 → CaM_N_0_C_1 + Ca | koff_cam_NT |
| CaM_N_1_C_1 → CaM_N_1_C_0 + Ca | koff_cam_CT |
| CaM_N_2_C_0 → CaM_N_1_C_0 + Ca | 2*koff_cam_NR |
| CaM_N_0_C_2 → CaM_N_0_C_1 + Ca | 2*koff_cam_CR |
| CaM_N_2_C_1 → CaM_N_1_C_1 + Ca | 2*koff_cam_NR |
| CaM_N_2_C_1 → CaM_N_2_C_0 + Ca | koff_cam_CT |
| CaM_N_1_C_2 → CaM_N_0_C_2 + Ca | koff_cam_NT |
| CaM_N_1_C_2 → CaM_N_1_C_1 + Ca | 2*koff_cam_CR |
| CaM_N_2_C_2 → CaM_N_1_C_2 + Ca | 2*koff_cam_NR |
| CaM_N_2_C_2 → CaM_N_2_C_1 + Ca | 2*koff_cam_CR |
| **Calcineurin Model.** | |
| CaN + CaM_N_2_C_2 ↔ CaN_CaM | kon_can_cam<br>koff_can_cam |
| **Calcinueurin Dephosphorylation of Dynamin** | |
| CaN_CaM + DYNp ↔ CaN_DYNp | kon_can_dyn<br>koff_can_dyn |
| CaN_DYNp → CaN_CaM, DYN | kcat_can |
| **Binding of dynamin to syndapin 1 (SYN1)** | |
| DYN + SYN1 ↔ DYN_SYN1 | kon_dyn_syn<br>koff_dyn_syn |
| **CDK5 phosphorylation of dynamin** | |
| DYN + CDK5 ↔ DYN_CDK5 | kon_cdk_dyn<br>koff_cdk_dyn |
| DYN_CDK5 ↔ DYNp + CDK5 | kcat_cdk |
| **Reclustering of vesicle proteins in rafts (pits)** | |
| SYB(membrane) + AP180 → SYB(raft) | kon_syb_ap180 |
| Syt(membrane) + AP2 → Syt(raft) | kon_syt_ap2 |
| **PKA/AC Model.** | |
| CaM_N_2_C_2 + AC18 ↔ AC18_CaM | kon_ac_cam<br>koff_ac_cam |
| AC18_CaM → AC18_CaM + cAMP | kcat_ac18 |
| cAMP → null | kdeg_camp |
| R2C2 + cAMP ↔ R2C2_cAMP | kon_r2c2_camp1<br>koff_r2c2_camp1 |
| R2C2_cAMP + cAMP ↔ R2C2_2cAMPbb | kon_r2c2_camp2bb<br>koff_r2c2_camp2bb |
| R2C2_cAMP + cAMP ↔ R2C2_2cAMPab | kon_r2c2_camp2ab<br>koff_r2c2_camp2ab |
| R2C2_2cAMPbb + cAMP ↔ R2C2_3cAMP | kon_r2c2_camp3bb<br>koff_r2c2_camp3bb |

| | |
|---|---|
| R2C2_2cAMPab + cAMP ↔ R2C2_3cAMP | kon_r2c2_camp3ab<br>koff_r2c2_camp3ab |
| R2C2_3cAMP + cAMP ↔ R2C2_4cAMP | kon_r2c2_camp4<br>koff_r2c2_camp4 |
| R2C2_2cAMPab ↔ R2C_2cAMPab + PKA | kact_pka_23<br>kdeact_pka_23 |
| R2C2_3cAMP ↔ R2C_3cAMP + PKA | kact_pka_23<br>kdeact_pka_23 |
| R2C2_4cAMP ↔ R2C_4cAMP + PKA | kact_pka_4<br>kdeact_pka_4 |
| R2C_2cAMPab + cAMP ↔ R2C_3cAMP | kon_r2c_camp3<br>koff_r2c_camp3 |
| R2C_3cAMP + cAMP ↔ R2C_4cAMP | kon_r2c_camp4<br>koff_r2c_camp4 |
| R2C_4cAMP ↔ R2_4cAMP + PKA | kact_pka_r2c<br>kdeact_pka_r2c |
| **(De)phosphorylation of synapsin.** | |
| PKA + synapsin ↔ PKA_synapsin | kon_PKA_syn<br>koff_PKA_syn |
| PKA_synapsin → PKA + synapsin_p | kcat_PKA_syn |
| PP2A + synapsin_p ↔ PP2A_synapsin_p | kon_pp2a_synapsin<br>koff_pp2a_synapsin |
| PP2A_synapsin_p → PP2A + synapsin | kcat_pp2a_syn |
| **Vesicle Clustering Model.** | |
| **Synapsin binding to actin(immobile)** | |
| Synapsin + actin ↔ Synapsin_actin | kon_synapsin_actin<br>koff_synapsin_actin |
| Synapsin_p + actin ↔ Synapsin_p_actin | kon_synapsin_actin<br>koff_synapsin_actin_p |
| **Tomosyn (de)phosphorylation** | |
| TOMO + CDK5 ↔ TOMO_CDK5 | kon_cdk_tomo<br>koff_cdk_tomo |
| TOMO_CDK5 → TOMO_p + CDK5 | kcat_cdk_tomo |
| TOMO_p + CaN_CaM ↔ TOMO_p_CaN_CaM | kon_can_tomo<br>koff_can_tomo |
| TOMO_p_CaN_CaM → TOMO + CaN_CaM | kcat_can_tomo |
| **Tomosyn binding Rab3** | |
| TOMO_p + Rab3 ↔ Rab3_TOMO_p | kon_rab3_TOMO_p<br>koff_rab3_TOMO_p |
| Rab3_TOMO →Rab3 + TOMO | koff_rab3_TOMO_p |
| **Rab3_TOMO binds synapsin.** | |
| Rab3_TOMO_p + synapsin ↔<br>Rab3_TOMO_p_synapsin | kon_TOMO_p_synapsin<br>koff_TOMO_p_synapsin |
| Rab3_TOMO + synapsin ↔ Rab3_TOMO_synapsin | kon_TOMO_synapsin<br>koff_TOMO_synapsin |
| **Synapsin Dimerisation (vesicle crosslinking)** | |

| | |
|---|---|
| Synapsin + synapsin ↔ synapsin_dimer | kon_synapsin_dimer<br>koff_synapsin_dimer |
| synapsin_dimer_p ↔ synapsin + synapsin_p | kon_synapsin_dimer<br>koff_synapsin_dimer_p |
| **Binding of proteins to vesicles (buffering)** | |
| **Synapsin:** | |
| ves_syn1_site + synapsin(cytosol) ↔ synapsin(ves) | kon_synapsin_ves<br>koff_synapsin_ves |
| ves_syn1_site + synapsin_p(cytosol) ↔ synapsin_p(ves) | kon_synapsin_ves<br>koff_synapsin_p_ves |
| **Complexin:** | |
| ves_CXN_site + CXN(cytosol) ↔ CXN(ves) | kon_cxn_ves<br>koff_cxn_ves |
| **Rab3-GTP:** | |
| ves_rab3_site + Rab3(cytosol) ↔ Rab3(ves) | kon_rab3_ves<br>koff_rab3_ves |
| **Munc13/18:** | |
| ves_m13_site + M13(cytosol) ↔ M13(ves) | kon_m13_ves<br>koff_m13_ves |
| ves_m18_site + M13(cytosol) ↔ M13(ves) | kon_m18_ves<br>koff_m18_ves |
| **Syndapin-1:** | |
| ves_syndap_site + SYN1(cyt) ↔ SYN(ves) | kon_syn1_ves<br>koff_syn1_ves |
| **AlphaSNAP:** | |
| ves_asnap_site + aSNAP(cyt) ↔ aSNAP | kon_asnap_ves<br>koff_asnap_ves |
| **NSF:** | |
| ves_nsf_site + NSF(cyt) ↔ NSF(ves) | kon_nsf_ves<br>koff_nsf_ves |

### References.


Althof, D., D. Baehrens, M. Watanabe, N. Suzuki, B. Fakler and Á. Kulik (2015). "Inhibitory and excitatory axon terminals share a common nano-architecture of their Cav2.1 (P/Q-type) Ca(2+) channels." Front Cell Neurosci **9**: 315.
Anggono, V., K. J. Smillie, M. E. Graham, V. A. Valova, M. A. Cousin and P. J. Robinson (2006). "Syndapin I is the phosphorylation-regulated dynamin I partner in synaptic vesicle endocytosis." Nat Neurosci **9**(6): 752-760.
Antunes, G. and E. De Schutter (2012). "A Stochastic Signaling Network Mediates the Probabilistic Induction of Cerebellar Long-Term Depression." Journal of Neuroscience **32**(27): 9288-9300.
Barak, B., A. Williams, N. Bielopolski, I. Gottfried, E. Okun, M. A. Brown, U. Matti, J. Rettig, E. L. Stuenkel and U. Ashery (2010). "Tomosyn expression pattern in the mouse hippocampus suggests both presynaptic and postsynaptic functions." Front Neuroanat **4**: 149.

Bartol, T. M., D. X. Keller, J. P. Kinney, C. L. Bajaj, K. M. Harris, T. J. Sejnowski and M. B. Kennedy (2015). "Computational reconstitution of spine calcium transients from individual proteins." Front Synaptic Neurosci **7**: 17.
Bhalla, U. S. and R. Iyengar (1999). "Emergent properties of networks of biological signaling pathways." Science **283**(5400): 381-387.
Bowen, M. E., K. Weninger, J. Ernst, S. Chu and A. T. Brunger (2005). "Single-molecule studies of synaptotagmin and complexin binding to the SNARE complex." Biophysical Journal **89**(1): 690-702.
Burkhardt, P., D. A. Hattendorf, W. I. Weis and D. Fasshauer (2008). "Munc18a controls SNARE assembly through its interaction with the syntaxin N-peptide." EMBO J **27**(7): 923-933.
Buxbaum, J. D. and Y. Dudai (1989). "A quantitative model for the kinetics of cAMP-dependent protein kinase (type II) activity. Long-term activation of the kinase and its possible relevance to learning and memory." J Biol Chem **264**(16): 9344-9351.
Chugh, J., A. Chatterjee, A. Kumar, R. K. Mishra, R. Mittal and R. V. Hosur (2006). "Structural characterization of the large soluble oligomers of the GTPase effector domain of dynamin." FEBS J **273**(2): 388-397.
Cipriano, D. J., J. Jung, S. Vivona, T. D. Fenn, A. T. Brunger and Z. Bryant (2013). "Processive ATP-driven Substrate Disassembly by the N-Ethylmaleimide-sensitive Factor (NSF) Molecular Machine." Journal of Biological Chemistry **288**(32): 23436-23445.
Coppola, T., S. Magnin-Luthi, V. Perret-Menoud, S. Gattesco, G. Schiavo and R. Regazzi (2001). "Direct interaction of the Rab3 effector RIM with Ca2+ channels, SNAP-25, and synaptotagmin." Journal of Biological Chemistry **276**(35): 32756-32762.
Davis, A. F., J. Bai, D. Fasshauer, M. J. Wolowick, J. L. Lewis and E. R. Chapman (1999). "Kinetics of synaptotagmin responses to Ca2+ and assembly with the core SNARE complex onto membranes." Neuron **24**(2): 363-376.
Diril, M. K., M. Wienisch, N. Jung, J. Klingauf and V. Haucke (2006). "Stonin 2 is an AP-2-dependent endocytic sorting adaptor for synaptotagmin internalization and recycling." Dev Cell **10**(2): 233-244.
Doi, T., S. Kuroda, T. Michikawa and M. Kawato (2005). "Inositol 1,4,5-trisphosphate-dependent Ca2+ threshold dynamics detect spike timing in cerebellar Purkinje cells." Journal of Neuroscience **25**(4): 950-961.
Faas, G. C., S. Raghavachari, J. E. Lisman and I. Mody (2011). "Calmodulin as a direct detector of Ca2+ signals." Nat Neurosci **14**(3): 301-304.
Faas, G. C., B. Schwaller, J. L. Vergara and I. Mody (2007). "Resolving the fast kinetics of cooperative binding: Ca2+ buffering by calretinin." PLoS Biol **5**(11): e311.
Fujita, Y., T. Sasaki, K. Fukui, H. Kotani, T. Kimura, Y. Hata, T. C. Sudhof, R. H. Scheller and Y. Takai (1996). "Phosphorylation of Munc-18/n-Sec1/rbSec1 by protein kinase C - Its implication in regulating the interaction of Munc-18/n-Sec1/rbSec1 with syntaxin." Journal of Biological Chemistry **271**(13): 7265-7268.
Fujita, Y., H. Shirataki, T. Sakisaka, T. Asakura, T. Ohya, H. Kotani, S. Yokoyama, H. Nishioka, Y. Matsuura, A. Mizoguchi, R. H. Scheller and Y. Takai (1998). "Tomosyn: a syntaxin-1-binding protein that forms a novel complex in the neurotransmitter release process." Neuron **20**(5): 905-915.
Gallimore, A. R., T. Kim, K. Tanaka-Yamamoto and E. De Schutter (2018). "Switching On Depression and Potentiation in the Cerebellum." Cell Reports **22**(3): 722-733.

Gordon, S. L. and M. A. Cousin (2014). "The Sybtraps: control of synaptobrevin traffic by synaptophysin, α-synuclein and AP-180." Traffic **15**(3): 245-254.
Hui, E., J. Bai, P. Wang, M. Sugimori, R. R. Llinas and E. R. Chapman (2005). "Three distinct kinetic groupings of the synaptotagmin family: candidate sensors for rapid and delayed exocytosis." Proc Natl Acad Sci U S A **102**(14): 5210-5214.
Imig, C., S. W. Min, S. Krinner, M. Arancillo, C. Rosenmund, T. C. Südhof, J. Rhee, N. Brose and B. H. Cooper (2014). "The morphological and molecular nature of synaptic vesicle priming at presynaptic active zones." Neuron **84**(2): 416-431.
Jimah, J. R. and J. E. Hinshaw (2019). "Structural Insights into the Mechanism of Dynamin Superfamily Proteins." Trends Cell Biol **29**(3): 257-273.
Kawaguchi, S. and T. Hirano (2013). "Gating of long-term depression by Ca2+/calmodulin-dependent protein kinaseII through enhanced cGMP signalling in cerebellar Purkinje cells." Journal of Physiology-London **591**(7): 1707-1730.
Kitagawa, Y., T. Hirano and S. Y. Kawaguchi (2009). "Prediction and validation of a mechanism to control the threshold for inhibitory synaptic plasticity." Molecular Systems Biology **5**: 16.
Lai, Y., U. B. Choi, J. Leitz, H. J. Rhee, C. Lee, B. Altas, M. L. Zhao, R. A. Pfuetzner, A. L. Wang, N. Brose, J. Rhee and A. T. Brunger (2017). "Molecular Mechanisms of Synaptic Vesicle Priming by Munc13 and Munc18." Neuron **95**(3): 591-+.
Li, Y. L., G. J. Augustine and K. Weninger (2007). "Kinetics of complexin binding to the SNARE complex: Correcting single molecule FRET measurements for hidden events." Biophysical Journal **93**(6): 2178-2187.
Liu, M., E. Girma, M. A. Glicksman and R. L. Stein (2010). "Kinetic mechanistic studies of Cdk5/p25-catalyzed H1P phosphorylation: metal effect and solvent kinetic isotope effect." Biochemistry **49**(23): 4921-4929.
Mezer, A., E. Nachliel, M. Gutman and U. Ashery (2004). "A new platform to study the molecular mechanisms of exocytosis." J Neurosci **24**(40): 8838-8846.
Millet, O., P. Bernadó, J. Garcia, J. Rizo and M. Pons (2002). "NMR measurement of the off rate from the first calcium-binding site of the synaptotagmin I C2A domain." FEBS Lett **516**(1-3): 93-96.
Murthy, V. N. and C. F. Stevens (1999). "Reversal of synaptic vesicle docking at central synapses." Nature Neuroscience **2**(6): 503-507.
Reshetniak, S., J. E. Ußling, E. Perego, B. Rammner, T. Schikorski, E. F. Fornasiero, S. Truckenbrodt, S. Köster and S. O. Rizzoli (2020). "A comparative analysis of the mobility of 45 proteins in the synaptic bouton." EMBO J **39**(16): e104596.
Rickman, C. and B. Davletov (2003). "Mechanism of calcium-independent synaptotagmin binding to target SNAREs." Journal of Biological Chemistry **278**(8): 5501-5504.
Ross, J. A., M. A. Digman, L. Wang, E. Gratton, J. P. Albanesi and D. M. Jameson (2011). "Oligomerization state of dynamin 2 in cell membranes using TIRF and number and brightness analysis." Biophys J **100**(3): L15-L17.
Shin, B. N., D. W. Kim, I. H. Kim, J. H. Park, J. H. Ahn, I. J. Kang, Y. L. Lee, C. H. Lee, I. K. Hwang, Y. M. Kim, S. Ryoo, T. K. Lee, M. H. Won and J. C. Lee (2019). "Down-regulation of cyclin-dependent kinase 5 attenuates p53-dependent apoptosis of hippocampal CA1 pyramidal neurons following transient cerebral ischemia." Sci Rep **9**(1): 13032.
Shu, T., H. Jin, J. E. Rothman and Y. Zhang (2020). "Munc13-1 MUN domain and Munc18-1 cooperatively chaperone SNARE assembly through a tetrameric complex." Proc Natl Acad Sci U S A **117**(2): 1036-1041.

Smillie, K. J. and M. A. Cousin (2005). "Dynamin I phosphorylation and the control of synaptic vesicle endocytosis." Biochem Soc Symp(72): 87-97.
Sun, T., X. S. Wu, J. Xu, B. D. McNeil, Z. P. Pang, W. Yang, L. Bai, S. Qadri, J. D. Molkentin, D. T. Yue and L. G. Wu (2010). "The role of calcium/calmodulin-activated calcineurin in rapid and slow endocytosis at central synapses." J Neurosci **30**(35): 11838-11847.
Takamori, S., M. Holt, K. Stenius, E. A. Lemke, M. Gronborg, D. Riedel, H. Urlaub, S. Schenck, B. Brugger, P. Ringler, S. A. Muller, B. Rammner, F. Grater, J. S. Hub, B. L. De Groot, G. Mieskes, Y. Moriyama, J. Klingauf, H. Grubmuller, J. Heuser, F. Wieland and R. Jahn (2006). "Molecular anatomy of a trafficking organelle." Cell **127**(4): 831-846.
Vivona, S., D. J. Cipriano, S. O'Leary, Y. H. Li, T. D. Fenn and A. T. Brunger (2013). "Disassembly of all SNARE complexes by N-ethylmaleimide-sensitive factor (NSF) is initiated by a conserved 1:1 interaction between α-soluble NSF attachment protein (SNAP) and SNARE complex." J Biol Chem **288**(34): 24984-24991.
Walter, A. M., K. Wiederhold, D. Bruns, D. Fasshauer and J. B. Sørensen (2010). "Synaptobrevin N-terminally bound to syntaxin-SNAP-25 defines the primed vesicle state in regulated exocytosis." J Cell Biol **188**(3): 401-413.
Wang, H. and D. R. Storm (2003). "Calmodulin-regulated adenylyl cyclases: cross-talk and plasticity in the central nervous system." Mol Pharmacol **63**(3): 463-468.
Wilhelm, B. G., S. Mandad, S. Truckenbrodt, K. Krohnert, C. Schafer, B. Rammner, S. J. Koo, G. A. Classen, M. Krauss, V. Haucke, H. Urlaub and S. O. Rizzoli (2014). "Composition of isolated synaptic boutons reveals the amounts of vesicle trafficking proteins." Science **344**(6187): 1023-1028.
Wu, Z., S. T. Wong and D. R. Storms (1993). "Modification of the calcium and calmodulin sensitivity of the type I adenylyl cyclase by mutagenesis of its calmodulin binding domain." J Biol Chem **268**(32): 23766-23768.
Zdanowicz, R., A. Kreutzberger, B. Liang, V. Kiessling, L. K. Tamm and D. S. Cafiso (2017). "Complexin Binding to Membranes and Acceptor t-SNAREs Explains Its Clamping Effect on Fusion." Biophys J **113**(6): 1235-1250.
Zikich, D., A. Mezer, F. Varoqueaux, A. Sheinin, H. J. Junge, E. Nachliel, R. Melamed, N. Brose, M. Gutman and U. Ashery (2008). "Vesicle priming and recruitment by ubMunc13-2 are differentially regulated by calcium and calmodulin." J Neurosci **28**(8): 1949-1960.
